# Genetically encoded tool for manipulation of ATP/ADP ratio in human cells

**DOI:** 10.1101/2025.08.12.670003

**Authors:** Alex E. Ekvik, Megan M. Kober, Denis V. Titov

## Abstract

The ability of cells to power energy-demanding processes depends on maintaining the ATP hydrolysis reaction a billion-fold away from equilibrium. Cells respond to changes in energy state by sensing changes in ATP, ADP, AMP, and inorganic phosphate. A key barrier to a better understanding of the maintenance of energy homeostasis is a lack of tools for direct manipulation of energy state in living cells. Here, we report the development of ATPGobble–a genetically encoded tool for controlling cellular ATP hydrolysis rate. We validated ATPGobble by showing that it doubles the energy demand, decreases [ATP]/[ADP] and [ATP]/[AMP] ratios, and activates AMPK activity in human cells. We then used ATPGobble to systematically characterize the proteome and phosphoproteome changes caused by direct manipulation of the energy state. Our results establish ATPGobble as a powerful approach for dissecting the regulatory roles of energy state in human cells, opening new opportunities to study how cellular energy state governs physiology, stress responses, and disease processes.

## INTRODUCTION

The ability of ATP to fuel energy-requiring reactions relies on the fact that the physiological ratio between ATP and its hydrolysis products, ADP and phosphate, is a billion-fold away from the reaction equilibrium, strongly favoring hydrolysis. Cells have a host of pathways and regulatory mechanisms that adjust ATP production and consumption rates to keep [ATP]/([ADP][Pi]) ratio far from equilibrium. Changes in energy homeostasis have been observed in multiple diseases (Ohta et al., 2017; Vishnu et al., 2021) and with interventions such as caloric restriction and exercise that have many beneficial effects. However, the causal relationships between changes in energy state and physiological effects have been difficult to establish due to a lack of tools for direct manipulation of cellular energy state.

Multiple mechanisms allow regulation of ATP production and consumption in response to changes in ATP, ADP, AMP and phosphate. Enzymes in energy producing pathways such as glycolysis (Lowry & Passonneau, 1966; Ureta, 1976; Weber, 1969) and the tricarboxylic acid cycle (Denton et al., 1975; Gabriel et al., 1985; Lawlis & Roche, 1981), are allosterically inhibited by ATP, while being allosterically activated by ADP, AMP and phosphate while enzyme in energy-demanding pathways such as gluconeogenesis (Sakai et al., 1987), and *de novo* biosynthesis of nucleotides (Caskey et al., 1964) have the opposite regulation. In addition to allostery, the relative changes in ATP, ADP and phosphate regulate energy metabolism through mass action as they are the substrates and products of enzymes in ATP producing and ATP consuming pathways. Beyond direct effects on metabolic enzymes, cellular energy state controls intracellular processes through posttranslational modifications where low ATP and high ADP, AMP and phosphate activate the AMP-sensitive kinase (AMPK) (Carling et al., 1987) and inhibit pyruvate dehydrogenase kinases (PDK1-4) (Biondi et al., 2002; Schulze et al., 2016a), which phosphorylate many targets and help maintain cellular energy homeostasis (Winder & Hardie, 1996). Finally, ATP/ADP ratio can control ATP-sensitive potassium channels and affect plasma membrane potential. The latter mechanism couples insulin secretion to glucose concentration in pancreatic β-cells through glucose-mediated increases in ATP/ADP ratio (Ashcroft & Rorsman, 1989). Tools for direct manipulation of cellular energy state would allow researchers to better understand how these regulatory mechanisms work together to regulate cellular energy state and potentially lead to discovery of new modes of regulation.

Current approaches to perturbing the cellular energy state include inhibition of ATP-producing pathways using small molecule inhibitors (Ibrahim et al., 2020) or genetic manipulations. These interventions are not specific to ATP/ADP ratio as their inhibition causes redox imbalance and prevents biosynthesis of most intracellular metabolites. Further, it is challenging to target drugs to specific tissues or subcellular compartments. Recently, genetically encoded tools for manipulation of metabolism (GEMMs) have been developed that use enzymes to directly modify the metabolic parameters such as NADH/NAD+, mitochondrial membrane potential, NADPH/NADP+, and CoQH2/CoQ in a compartment-specific manner (Choe & Titov, 2022).

Here, we introduce ATPGobble – a GEMM that decreases ATP/ADP ratio in human cells. To identify promising GEMM candidates for manipulating ATP/ADP ratio, we screened ATPases for their activity in human cells and identified *E. coli* F1 ATPase as the most promising candidate, which we called ATPGobble. We have validated ATPGobble by demonstrating that it doubles the energy demand of the cell, decreases ATP/ADP and ATP/AMP ratios, and activates AMPK kinase, consistent with expected activities of a GEMM that lowers ATP/ADP ratio. We then used ATPGobble to systematically profile the responses of human cells to direct manipulation of energy state using proteomics and phosphoproteomics. The proteomic analysis revealed concerted activation of ATP-producing pathways through the upregulation of glucose transporters and pyruvate dehydrogenase (PDH) activity, and downregulation of ATP-consuming pathways, such as translation, fatty acid biosynthesis, and cell growth. We also identified and validated changes in proteins not previously associated with the energy state, such as the Olduvai domain-containing NBPF protein family, which has expanded a proposed role in human-specific cognition (Glunčić et al., 2024). Our results establish ATPGobble as a novel tool for studying the regulation of cellular energy in health and disease.

## RESULTS

### Screen of ATP-consuming enzymes identifies *E. coli* F1 ATPase as promising ATPGobble candidate

To identify a GEMM candidate that can directly lower the ATP/ADP ratio in living cells, we compiled a list of enzymes with ATPase activity and no other substrates based on the BRENDA database (Figure S1). Most of the candidates were extracellular proteins, mainly apyrases, which was expected, as expressing an intracellular ATPase with no other activity is unlikely to provide an evolutionary advantage. We removed signal peptides and transmembrane domains to ensure cytosolic localization of enzymes. We expressed FLAG-tagged versions of each candidate using a doxycycline-inducible promoter in HeLa cells infected with lentiviruses based on LVX-TetOne-Puro constructs and screened for ATPase activity. We confirmed the expression of the candidate proteins by a FLAG western blot (Figure S1B). To estimate the ATPase activity of the candidates, we performed a lactate assay to assess the effect of the constructs on the fermentation rate (Figures S1A). The rationale behind this experiment is based on the fact that fermentation rate is coupled to ATP demand of the cells; hence, ATPase activity would result in an increased lactate production. The expression of α, β and γ subunits of the *E. coli* F1 ATPase caused by far the largest increase in lactate secretion, indicating elevated ATP consumption (Figure S1A). F1 ATPase forms the catalytic part of ATP synthase and, when expressed independently from the proton channel, F0, can hydrolyze ATP instead of synthesizing it (Figure 1A). Overexpression of *E. coli* F1 ATPase has been shown to stimulate fermentation in bacterial cells (Boecker et al., 2021; Koebmann et al., 2002), further validating its promise as a GEMM. Therefore, we chose to focus on *E. coli* F1 ATPase, referred to as ATPGobble henceforth, as a genetically encoded tool for manipulation of cellular energy state.

**Figure 1.**
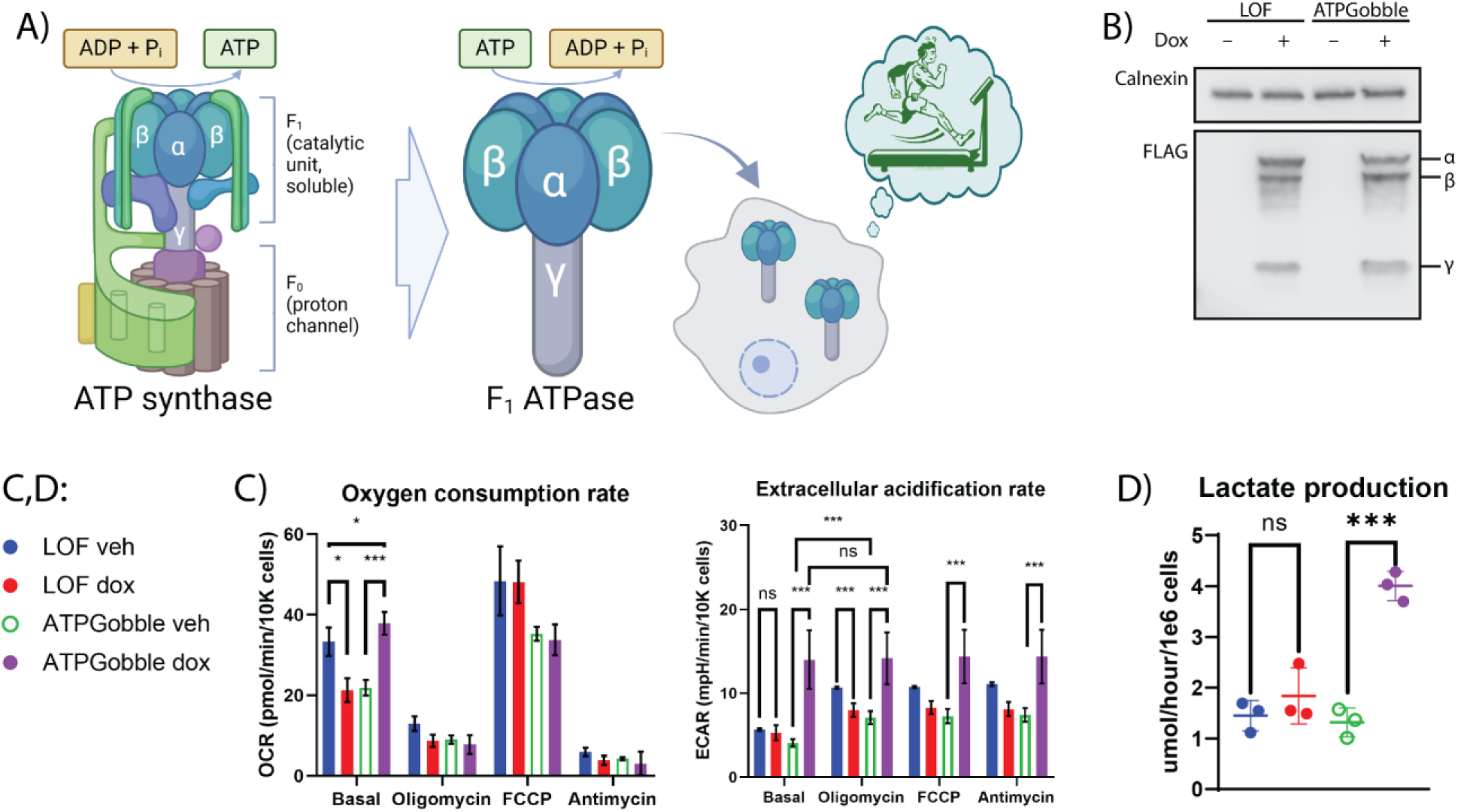
ATPGobble doubles the ATP production rate of human cells. A) The mechanism of the ATPGobble activity of *E. coli* F1 ATPase when used as ATPGobble. B) Doxycycline-inducible expression of ATPGobble subunits and the catalytically inactive loss-of-function (LOF) subunits (βK155Q; α and γ unaltered) in hTERT-RPE1 cells. Representative of 3 experiments. C) ATPGobble increases oxygen consumption rate (OCR) and extracellular acidification rate (ECAR) in hTERT-RPE1 cells. DMEM and water used as a vehicle. 3 experiments 5 wells per group each. Repeated measurements ANOVA followed by Fisher’s LSD test for main row effect with Tukey adjustment. D) ATPGobble increases lactate production in hTERT-RPE1 cells. 3 experiments 1 dish per group each. Paired homoscedastic t-test. ns > 0.05, * < 0.05, ** < 0.01 *** < 0.001. dox: doxycycline, veh: vehicle. Experiment means ± standard deviation of experiment means.

### ATPGobble doubles the ATP production rate of human cells

We next investigated the effect of ATPGobble expression on ATP-producing pathways. While the initial screening was done in HeLa cells (Figure S1), we performed most of our experiments in diploid human cell line hTERT-RPE1. To allow rigorously controlled experiments, we used a doxycycline-inducible promoter to acutely express ATPGobble and used a catalytically inactive mutant with a K155Q substitution in subunit β (Senior & al-Shawi, 1992), referred to as LOF (loss-of-function) henceforth, as a negative control. We verified that the expression of ATPGobble and LOF subunits in hTERT-RPE1 was adequate and doxycycline-dependent in both ATPGobble and LOF cell lines (Figure 1B).

We measured the effect of LOF and ATPGobble on the rate of ATP production by fermentation and respiration (Figure 1C,D). We found that LOF was expressed at the same level as ATPGobble (Figure 1B) and had no significant impact on the rates of ATP-producing pathways. On the other hand, ATPGobble expression increased fermentation rate 3-fold and respiration rate 2-fold (Figure 1C,D). Notably, the cells expressing ATPGobble showed no increase in their extracellular acidification in response to oligomycin, FCCP or antimycin, suggesting that ATPGobble saturated the glycolytic capacity of the cells.

### ATPGobble decreases ATP/ADP and ATP/AMP ratios and activates AMPK in human cells

We next tested the effect of ATPGobble on ATP/ADP and ATP/AMP ratios using LC-MS and PercevalHR–fluorescent sensor of ATP/ADP ratio (Tantama et al., 2013). We extracted metabolites from hTERT-RPE1 cells expressing ATPGobble and conducted targeted LC/MS of ATP, ADP and AMP (Figure 2A). ATPGobble expression decreased ATP/ADP ratio 2-fold and ATP/AMP ratio 3-fold (Figure 2A). A larger change in AMP is expected due to the conversion of ADP into ATP and AMP carried out by adenylate kinase (Atkinson, 1968). The change in ratio was mostly due to the increase in AMP and ADP, while ATP pools remained relatively stable (Figure S2). This was expected, as most of the adenine pool is in the form of ATP, and the conversion of ATP to ADP and AMP will result in larger relative changes of the latter. We confirmed the effect on the ATP/ADP ratio in live cells using a ATP/ADP ratio fluorescent sensor PercevalHR (Figure 2B). LOF had no effect on ATP/ADP or AMP/ADP ratio in any of the assays, confirming that changes observed using ATPGobble are due to its ATPase activity.

**Figure 2.**
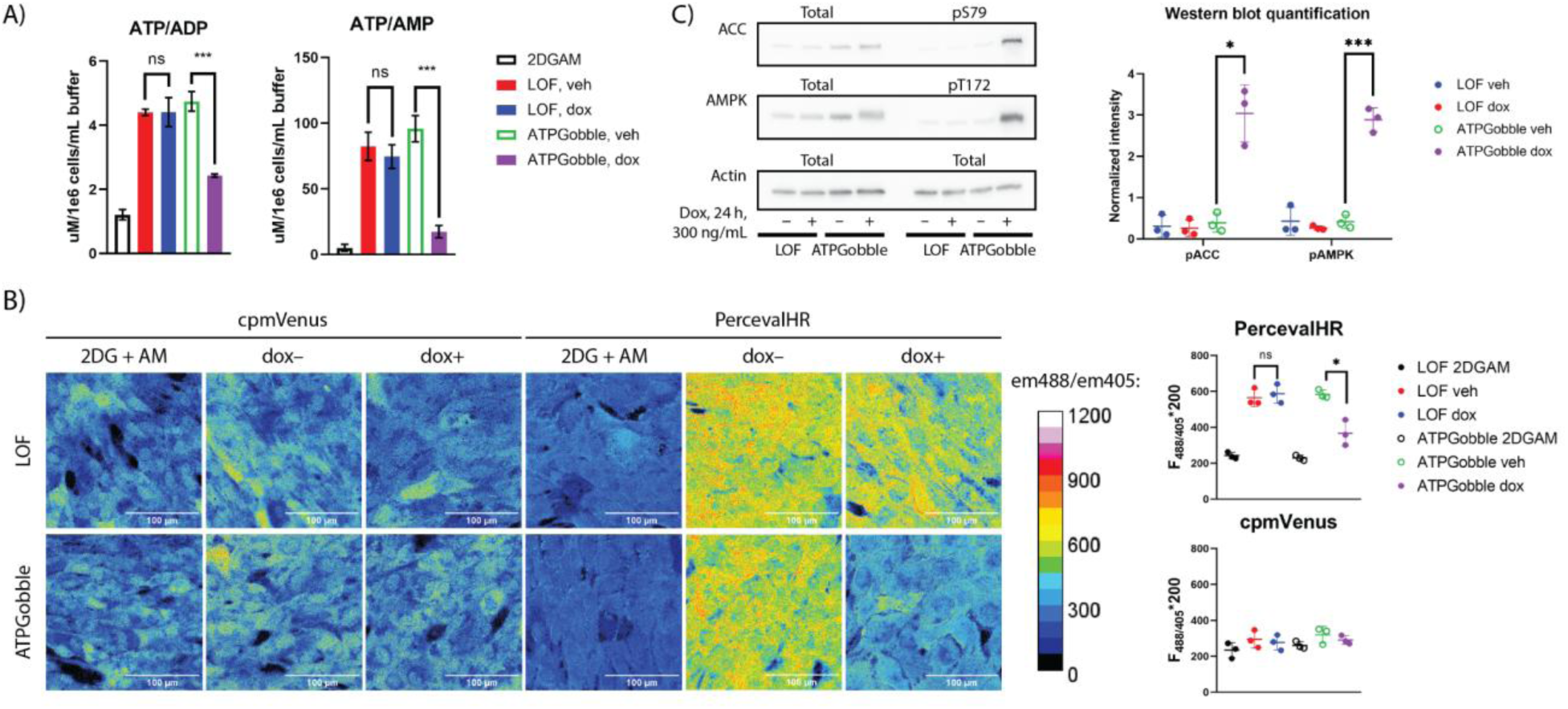
ATPGobble causes a decrease in ATP/ADP ratio in hTERT-RPE1 cells. A) LCMS quantification of ATP/ADP and ATP/AMP ratios after 24-hour ATPGobble induction. 3 experiments 5 dishes per group each. Repeated measurements ANOVA followed by Fisher’s LSD test for main row effect with Tukey adjustment. B) Pseudocolor images of PercevalHR and its non-ATP-sensitive control cpmVenus with and without ATPGobble induction. Em488/em405 is an indicator of ATP/ADP ratio; the original ratio was multiplied by 200 for easier visualization. 3 experiments, 1 well and 5 images each. Paired homoscedastic t-test. C) Phosphorylation of AMPKα and ACC in ATPGobble-expressing hTERT-RPE1 cells, a marker of AMPK activity. 3 experiments 1 lane per group each. Paired homoscedastic t-test. ns > 0.05, * < 0.05, ** < 0.01 *** < 0.001. dox: doxycycline, veh: vehicle. Experiment means ± standard deviation of experiment means.

We next asked if the decrease in ATP/AMP ratio observed with the induction of ATPGobble was enough to activate AMPK, by running a western blot of AMPK phosphorylation targets ACC S79 and AMPKα T172 (Rubink & Winder, 2005; Woods et al., 2003). In line with the decreased ATP/AMP ratio and as a result increased AMPK activity, AMPK targets ACC S79 and AMPK T172 were phosphorylated upon 24 hours of induction in the ATPGobble cells, but not in the LOF cells (Figure 2C).

Together, these results demonstrate that the expression of ATPGobble, but not LOF, causes large-scale perturbation of cellular energy metabolism. This validates ATPGobble as a genetically encoded tool for manipulation of energy homeostasis.

### ATPGobble causes a slowdown in cell proliferation rate

Having validated ATPGobble as a tool to manipulate cellular energy state, we investigated its effects on cell physiology. We reasoned that adaptations to the increase in cellular ATP demand due to the expression of ATPGobble may not only include the activation of ATP-producing pathway, as seen in Figure 1C&D, but also inhibition of ATP-consuming processes. Hence, we tested the effect of ATPGobble on cell growth and proliferation, a major energy-consuming process. We measured the proliferation rate of hTERT-RPE1 expressing LOF and ATPGobble in a doxycycline-inducible manner by Hoechst staining and subsequent nucleus counting, over the course of three days of induction or vehicle treatment (Figure 3A). Though all four treatment groups grew exponentially (Figure 3A), the expression of ATPGobble in hTERT-RPE1 cells led to an inhibition of growth rate by around 80%. Of note, the expression of LOF led to a 30% slowdown in growth, possibly due to the stress associated with the high levels of exogenous protein expression. We also asked if ATPGobble expression causes cell death by co-staining with SytoxGreen, a fluorescent DNA binding dye that preferentially stains cells with compromised plasma membranes. This revealed that SytoxGreen-positive cells made up less than 5% of the population in all groups, except for near 100% death in the positive control cells treated with 2-deoxyglucose and antimycin (Figure 3B).

**Figure 3.**
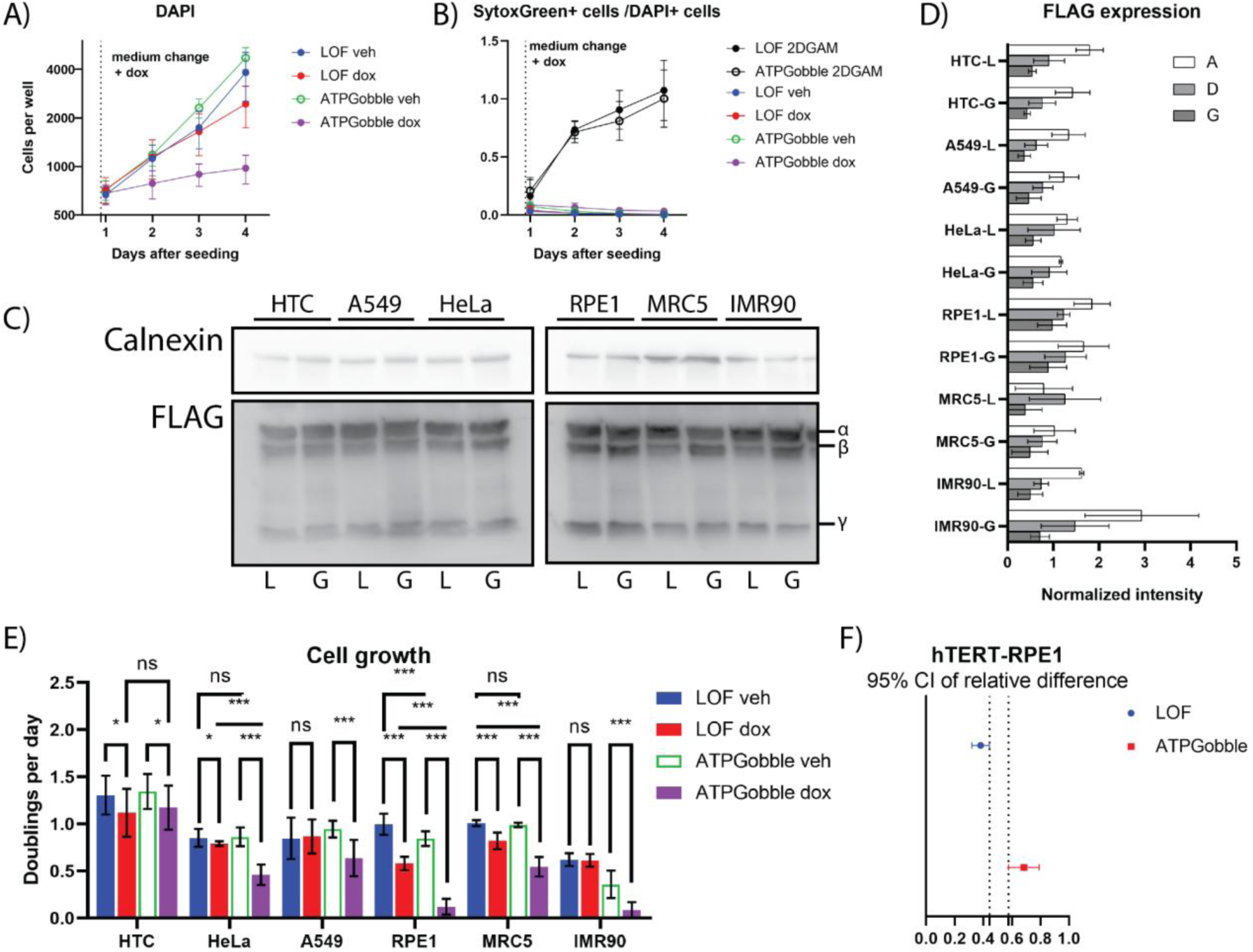
ATPGobble inhibits growth and translation. DAPI (A) and SytoxGreen (B) staining of RPE1 cells with 2-deoxyglucose and antimycin (2DGAM) as a positive control for SytoxGreen. 3 experiments with 8 cell wells per group in each experiment. C) Expression of ATPGobble and LOF subunits at 24 hours post-induction in the cell lines used for the cell growth assay. Representative of 3 experiments quantified in D). E) Doubling rates calculated from DAPI staining for different cell lines expressing ATPGobble or LOF in a dox-inducible manner. 3 experiments with 8 cell wells per treatment group in each experiment. Repeated measurements ANOVA followed by Fisher’s LSD test for main row effect with Tukey adjustment. F) 95% confidence intervals of the relative effect of induction on cell growth in hTERT-RPE1-LOF and hTERT-RPE1-ATPGobble ATPGobble cells, calculated from the data shown in E. ns > 0.05, * < 0.05, ** < 0.01 *** < 0.001. dox: doxycycline, veh: vehicle. Experiment means ± standard deviation of experiment means.

To systematically determine the effect of ATPGobble on cell proliferation rate, we measured six other transformed (HeLa, A549, HTC) and non-transformed cell lines (IMR90 amd MRC5). The effect of ATPGobble ranged from no effect in HTC cells to slowdown similar to hTERT-RPE1 in IMR90 cells (Figure 3D). The cell lines were generated side-by-side through lentiviral infection, including a replicate hTERT-RPE1 cell line. The expression of the ATPGobble subunits among the cell lines was comparable as verified by a FLAG western blot (Figure 3E&F), suggesting that different cell types may react differently to the manipulation of energy state.

### ATPGobble activity results in proteomic and phosphoproteomic changes

To investigate how cells adapt to direct manipulation of energy state, we measured the effect of 24-hour ATPGobble expression on total protein and phosphopeptide levels. The treatment groups were as follows: ATPGobble, vehicle; ATPGobble, doxycycline-induced; LOF, vehicle; LOF, doxycycline-induced. We used a Data Independent Acquisition (DIA) method on a timsTOF instrument, and the data was analyzed using DIA-NN (Demichev et al., 2020). After filtering and imputation, the total proteomics dataset contained 8419 proteins (Attachment 1) and the phosphoproteomics dataset contained 4357 entries (Attachment 2). In this section, we will describe the broad changes that we observed, and in the next two sections, we will focus on changes in energy-producing and energy-consuming pathways.

We observed a large-scale rearrangement of the proteome and phosphoproteome with 197 proteins (Figure 4A; Attachment 1) and 1152 (Figure 5A; Attachment 2) phosphopeptides changing more than 2-fold at the cutoff of 1% FDR Hochberg-Benjamini. 156 of the differentially expressed proteins were downregulated; for phosphopeptides, the split in the direction was more even with 576 of the phosphopeptides downregulated. Further, the PCA of the top highest-variance genes of the total (Figure 4C) and phosphoproteomics (Figure 5B) datasets showed a clear separation of the samples expressing ATPGobble. This confirmed that the vast majority of the changes in total protein and phosphopeptide levels were driven by ATPGobble and not LOF expression, both validating the LOF construct as a negative control and indicating that the ATP consumption by ATPGobble results in large-scale proteomic changes.

**Figure 4.**
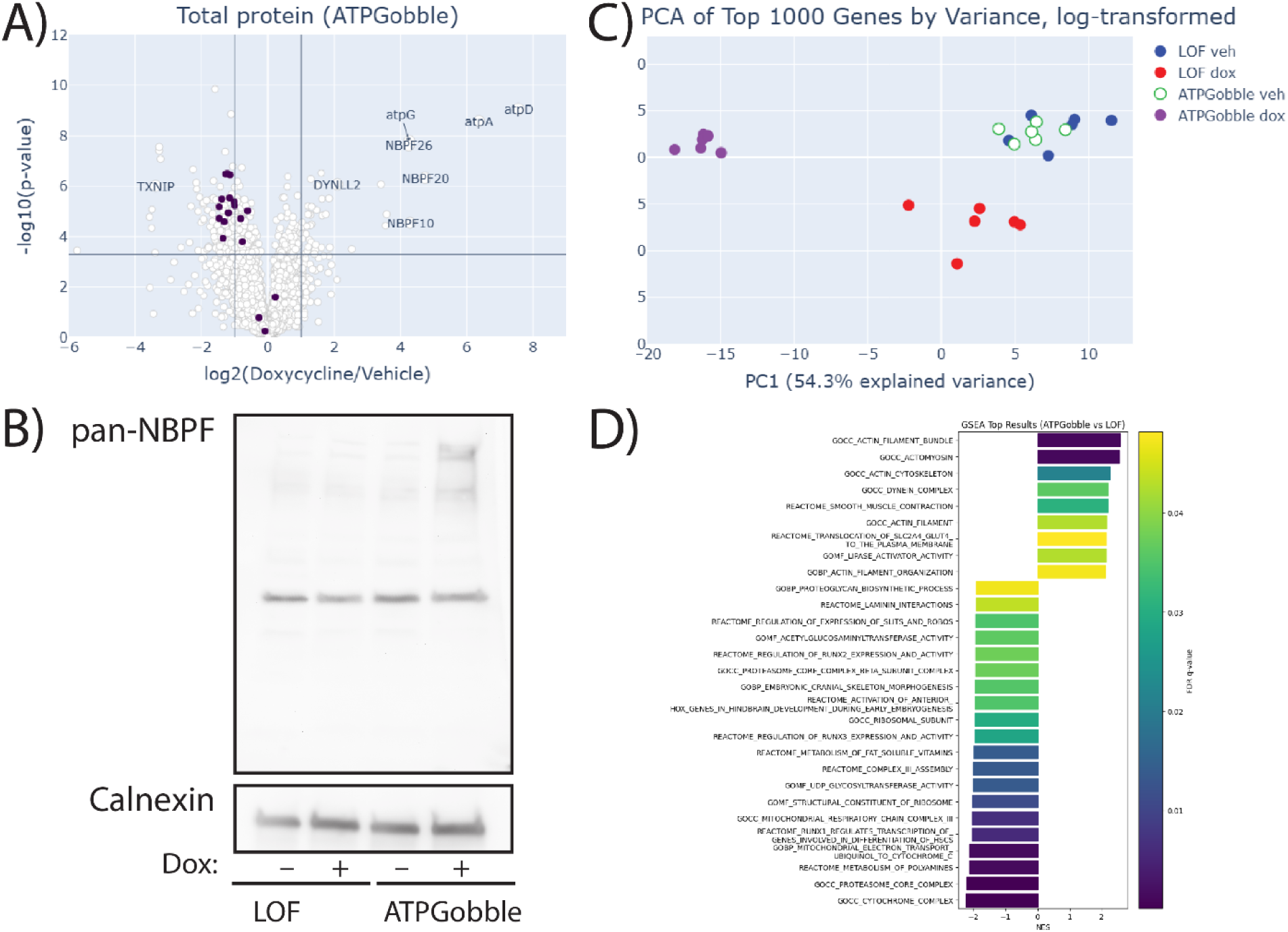
ATPGobble activity results in total protein changes in doxycycline-induced vs vehicle-treated RPE1-ATPGobble cells. A) Total protein expression in hTERT-RPE1-ATPGobble cells at 24 hours doxycycline vs vehicle treatment, with the members of GOCC Proteasome Core Complex highlighted in dark purple. Cutoff lines are 0.01% FDR (Hochberg-Benjamini) at log fold change of 1. P-values are from a two-tailed unpaired homoscedastic t-test. B) A western blot of NBPF family member expression in hTERT-RPE1-LOF cells and hTERT-RPE1-ATPGobble cells at 24 hours of doxycycline or vehicle treatment. Representative of 3 experiments. C) PCA of the total proteomics data set including the top 1000 genes by variance excluding ATPGobble genes, from hTERT-RPE1-LOF and hTERT-RPE1-ATPGobble cells treated with vehicle and doxycycline. D) Gene sets significantly enriched in GSEA at the FDR=5% cutoff, ATPGobble doxycycline vs LOF doxycycline. Negative NES signifies decreased expression in the ATPGobble doxycycline group.

**Figure 5.**
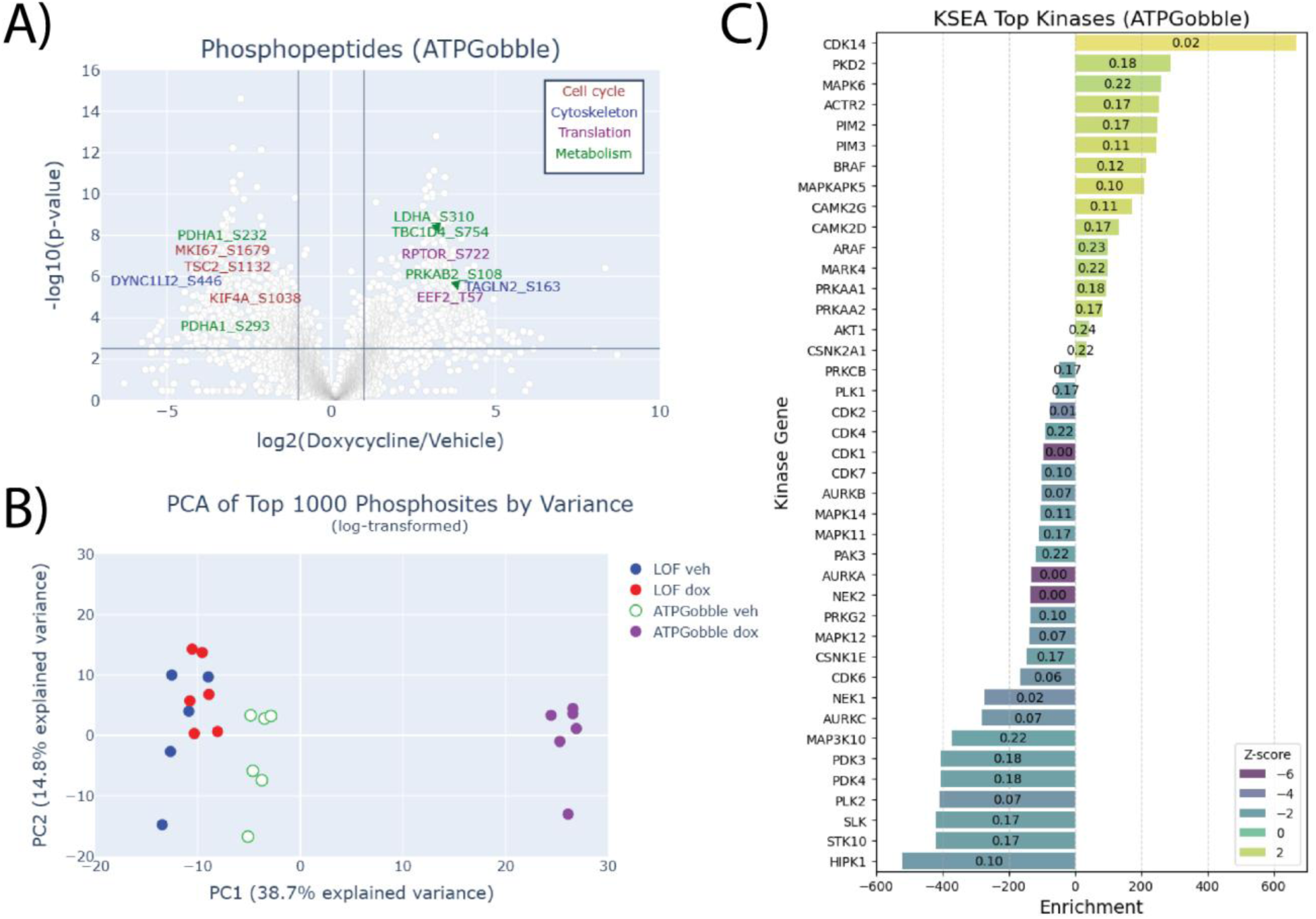
ATPGobble elicits changes in the phosphorylation of proteins regulating metabolism, translation, cell growth and cytoskeletal dynamics. A) Phosphoproteomic changes in hTERT-RPE1-ATPGobble cells after doxycycline induction vs vehicle treatment, with select examples of significant targets color-coded by their function. Cutoff lines are 1% FDR (Hochberg-Benjamini) at log fold change of 1. P-values are from a two-tailed unpaired homoscedastic t-test. B) PCA of the phosphoproteomic dataset including hTERT-RPE1-LOF and hTERT-RPE1-ATPGobble cells treated with vehicle or doxycycline, based on the top 1000 genes by variance. C) Significantly affected kinases detected by kinase set enrichment analysis at FDR < 0.25 and p-value < 0.05, colored by Z-score and labeled with FDR, in hTERT-RPE1-ATPGobble (D) cells.

On an individual protein level, the top upregulated proteins that were significant after multiple hypothesis testing correction were the ATPGobble subunits, and the top downregulated protein was TXNIP – a validated target of AMPK that is discussed in the next section. Beyond ATPGobble subunits, the most upregulated proteins were NBPF10, NBPF20 and NBPF26 (Figure 4A; Figure S3A&B; Attachment 1). NBPF26 was also phosphorylated at S1197 (Attachment 2). These proteins are members of the DUF1220/Olduvai domain-containing family with a poorly understood function that have been implicated in human brain development and experience a rapid human lineage-specific increase in copy number (Schulze et al., 2016b). We validated the upregulation of these proteins using a western blot with a pan-NBPF antibody, where we observed an induction of several bands, located around or above the 250 kDa marker upon ATPGobble expression, which corresponds to the predicted sizes of 435.6 kDa for NBPF10, and 646 kDa for NBPF20 and 190 kDa (Figure 4B). Together, this evidence suggests that the induction of ATPGobble leads to the upregulation of NBPF proteins.

We next performed Gene Set Enrichment Analysis (GSEA) to identify protein sets that might be regulated by the energy state of the cells but are not significant on single-protein level. GSEA revealed significant changes in both LOF and ATPGobble datasets (Attachment 3) which is why we have focused on the comparison between the two doxycycline-induced groups (Figure 4D). Overall, there were more downregulated gene sets than upregulated. The top-downregulated gene set was the proteasome core subunits (Figure 4A; Figure S3C&D). We observed a concerted downregulation by approximately 2-fold, which is reminiscent of proteasome disassembly with subsequent autophagy of the core particles, previously observed during nitrogen starvation in yeast (Waite et al., 2016). We also observed negative enrichment gene sets related to cell proliferation, extracellular protein synthesis, metabolism of fat-soluble vitamins and, somewhat surprisingly, cytochrome complex (Figure 4D). Positively enriched gene sets included cytoskeleton proteins, vesicle transport and relocation of the GLUT4 transporter as well as lipase activity.

We performed the kinase set enrichment analysis (KSEA) on phosphoproteomic data that revealed changes in the activity of kinases regulating metabolism, cell cycle and cytoskeletal dynamics (Figure 5C&D). As expected, AMPK subunits PRKAA2 and PRKAA1 were among positively enriched kinases, with its own subunits PRKAB1 (S96) and PRKAB2 (S108), as well as RPTOR (S722) among the top hyperphosphorylated targets (Attachment 5). Similarly, PDK isoforms PDK3 and PDK4 were significantly enriched with only one target, PDHA1. In addition, a range of cell cycle regulating kinases were affected in both ATPGobble and LOF cell lines.

In summary, we have generated a unique dataset that can serve as a signature of energy depletion in future GSEA studies.

### ATPGobble activates energy-producing pathways

As ATPGobble doubled the ATP production by hTERT-RPE1 cells (Figure 2), we investigated whether the activation of energy-producing pathways occurs in part at the proteomic or phosphoproteomic level. We first checked the expression of enzyme in energy producing pathways and were surprised to see no upregulation of enzymes in glycolysis, tricarboxylic acid cycle, and oxidative phosphorylation enzymes (Figure 6 D, Figure S5E&F), suggesting that the more than 2-fold activation of these pathways that we observed in Figure 1 is achieved through mass action, allosteric control, or posttranslational control of these pathways. In fact, the GSEA revealed multiple negatively enriched gene sets related to the mitochondrial complex III activity (Figure 4D).

**Figure 6.**
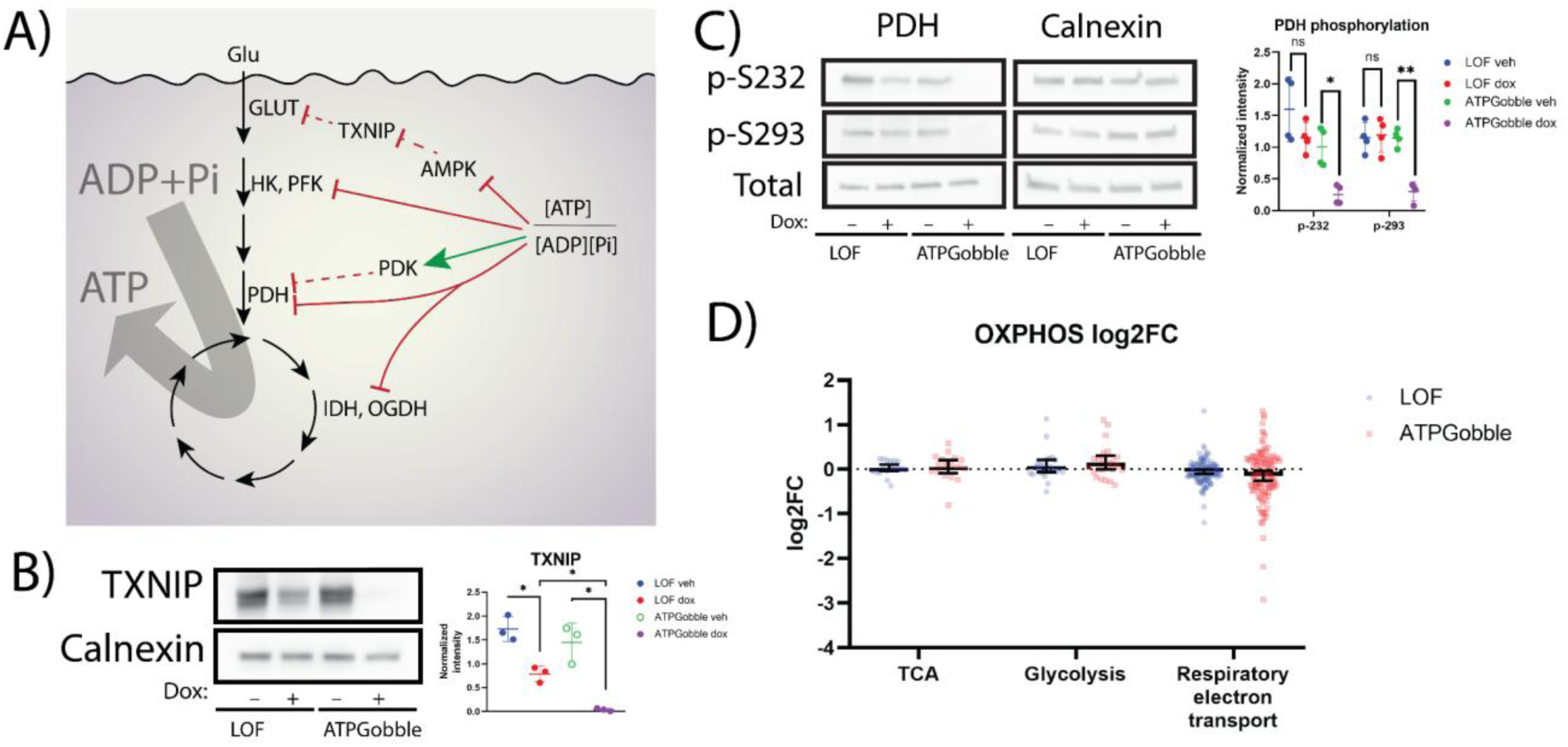
ATPGobble elicits changes in energy-producing processes. A) Summary of the effects of ATPGobble on energy-producing processes as observed in this paper. B) Decreased TXNIP expression in cells expressing ATPGobble. Representative of 3 experiments. C) Western blot revealing dephosphorylation of pyruvate dehydrogenase in ATPGobble-producing cells. Representative of 4 experiments. Significance in B and C was calculated using a paired homoscedastic t-test. D) 95% confidence intervals of the gene set mean log2 fold change in energy metabolism genes in hTERT-RPE1-ATPGobble and LOF cells after doxycycline vs vehicle treatment. ns > 0.05, * < 0.05, ** < 0.01 *** < 0.001. dox: doxycycline, veh: vehicle. Experiment means ± standard deviation of experiment means.

The energy producing pathways were posttranslationally activated upon the induction of ATPGobble, but not LOF, in the following ways. First, we saw several changes that would promote glucose transporter localization of GLUT1 and GLUT4 to the plasma membrane in the presence of ATPGobble, including TXNIP downregulation and phosphorylation of TBC1D4. TXNIP is an α-arrestin protein that promotes that clathrin-mediated endocytosis of GLUT1 and is phosphorylated by AMPK at Ser308, which leads to its degradation, hence promoting GLUT1 localization to the plasma membrane under conditions of energy stress (Wu et al., 2013). We have validated ATPGobble-driven downregulation of TXNIP by a western blot (Figure 6B). We could not detect TXNIP in our phosphoproteomics dataset to confirm whether it was phosphorylated. TBC1D4 is a Rab-GTPase-activating protein that regulates translocation of GLUT4 containing vesicles to the plasma membrane. We found that phosphorylation of TBC1D4 at S754 is increased nearly 10-fold in the presence of ATPGobble. In mouse, S761 (analogous position to human S754) has previously been shown to be phosphorylated by AMPK, which leads to dissociation of TBC1D4 from GLUT4-storage vesicles (Eickelschulte et al., 2021; Treebak et al., 2010). Further, we saw a positive enrichment of the GO code associated with GLUT4 transporter relocation (Figure 4C; Figure S5 A&B), as well as a pronounced increase in the phosphorylation of the proteins within the same gene set, including TBC1D4 at S754 (Figure S5 C&D). Second, we found that among the top negatively enriched kinases in our phosphoproteomics dataset were PDK3 and PDK4, which share the same target PDHA1 (S232 and S293) (Figure 5C). The phosphorylation inhibits PDHA1, limiting the influx into the tricarboxylic acid cycle and the activity of PDKs *in vitro* is well known to be directly regulated by ADP (Hucho et al., 1972; Pratt & Roche, 1979). We verified the dephosphorylation of PDH at both of these residues by a western blot (Figure 6C). Finally, our GSEA revealed an upregulation in genes responsible for lipase activity. In summary, ATPGobble increases the activity of fermentation and respiration pathways through a combination of mass action and allostery effect, induction of glucose transporters by downregulating TXNIP and phosphorylating TBC1D4, and activation of PDH activity through dephosphorylation (Figure 6A).

### ATPGobble inhibits energy-consuming processes

Finally, we investigated the effect of ATPGobble on energy-consuming processes. We have shown above that ATPGobble induces the phosphorylation of ACC by AMPK (Figure 2C), which inhibits *de novo* fatty acid biosynthesis, and here we investigated the effect of ATPGobble on other energy-consuming processes, such as translation, cytoskeleton dynamics, and proliferation.

Translation is downregulated in our dataset in a number of ways. We found that the translation elongation factor EEF2 was phosphorylated at T57 in the ATPGobble-expressing samples but none of the control samples. This was confirmed by a western blot (Figure 7B,C). This phosphorylation is known to inhibit translation and is carried out by EEF2 kinase (EEF2K), an AMPK target (Browne et al., 2004). Additionally, we saw multiple previously uncharacterized changes in the phosphorylation of translation initiation factors and ribosomal subunits (Attachment 1). Further, in line with the observed decrease in translation rates in ATPGobble-expressing cells, we found that genes within the GO MF Structural Constituent of Ribosome set were significantly downregulated, a gene set in which large subunit, small subunit and mitochondrial ribosome genes were all among the top downregulated genes (Figure S4 A&B). With this in mind, we tested if the rate of translation was inhibited upon the induction of ATPGobble: we metabolically labelled nascent peptides with azidohomoalanine and after click-chemistry labeling with biotin conducted a western blot. We found that the nascent peptide levels in the cells expressing ATPGobble were lower than in the controls, confirming that ATPGobble activity inhibits translation (Figure 7D,E). The downregulation of the proteasome levels in the presence of ATPGobble (Figure 4A, D; Figure S3 C&D) may further allow cells to slow down protein turnover that can be a significant source of energy demand even in the absence of growth.

**Figure 7.**
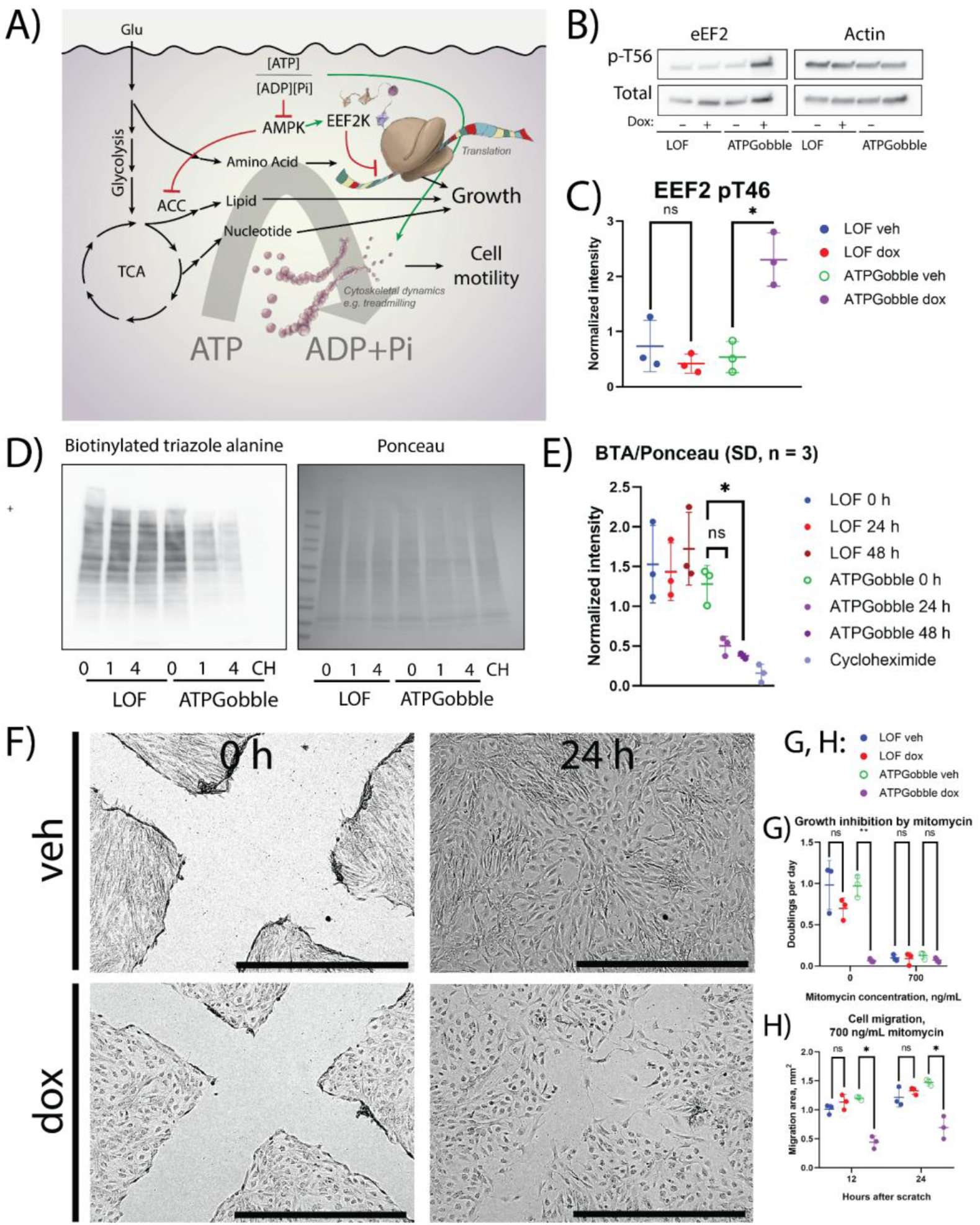
Effect of ATPGobble on energy-consuming processes. A) A schematic summary of energy-consuming processes changed by ATPGobble as observed in this paper. B) Phosphorylation of eEF2 in cells expressing ATPGobble and C) its quantification. D) Translation rates in RPE1-ATPGobble and RPE1-LOF cells. Nascent peptides were metabolically labelled by incubating the cells with azidohomoalanine for 4 hours. CH = cycloheximide and E) their quantification. F) Representative images of hTERT-ATPGobble cells migrating to the scratch area after doxycycline induction or vehicle treatment. G) Validation of the inhibition by mitomycin of the growth rate of hTERT-RPE1-LOF and hTERT-RPE1-ATPGobble cells. H) Quantification of migration area by the cells expressing ATPGobble and LOF after a scratch, growth inhibited by mitomycin. ns > 0.05, * < 0.05, ** < 0.01 *** < 0.001. dox: doxycycline, veh: vehicle. Experiment means ± standard deviation of experiment means.

Another energy-consuming process is cytoskeletal dynamics, such as polymerization as a part of cytoskeletal building blocks and cargo transport using microtubules. Our proteomics and phosphoproteomics datasets pointed to a shift in cytoskeletal dynamics on several levels. Our phosphoproteomics dataset revealed that cytoskeleton regulation proteins were among the top differentially phosphorylated targets. For example, while DYNLL2 was among the top differentially expressed targets in our proteomics dataset, and GSEA returned the GO CC Dynein Complex gene set as positively enriched, DYNC1L12 was also significantly hypophosphorylated (Figure 4A,D; Figure 5A). Because one of the main processes affected by cytoskeleton dynamics is migration, we asked if ATPGobble-expressing cells had an impaired migration ability through a series of scratch assays (Figure 7F–H). Mitomycin treatment abolished the differences in cell proliferation discussed earlier (Figure 7H); mitomycin was added at the same time as doxycycline. Even with these differences abolished, we found that ATPGobble-expressing cells had a significantly impaired migration performance at 12 and 24 hours after the scratch, which was in line with a previous report that energy stress inhibits migration. Our GSEA also revealed a positive enrichment of multiple gene sets associated with cytoskeleton (Figure 4D; Figure S6C–F); Interestingly, among the top upregulated kinases detected in KSEA was CDK14, its main target being TAGLN2 S163 (Figure 5A,C). CDK14 has been reported to promote cell migration by inhibiting TAGLN2 through this phosphorylation (Leung et al., 2011), and our observation in that light hints at a synergistic relationship between the ability of CDK14 to promote migration, and energy stress.

Finally, in line with the previously observed effects of ATPGobble on cell proliferation, our phosphoproteomics dataset showed changes in the regulation of cell cycle proteins. The cell cycle kinases enriched in KSEA included downregulation in multiple cyclin-dependent kinases, as well as a range of MAP kinases (Figure 5C). Notably, some of these kinases were also significantly downregulated in LOF cells, but the magnitude of these changes was much lower (Figure S4C).

## DISCUSSION

Here we developed and validated ATPGobble as the genetically encoded tool for manipulation of energy state in human cells. We showed that ATPGobble doubles the energy production and decreases the ATP/ADP ratio. We have also developed LOF–a catalytically dead version of ATPGobble that can serve as a control. Together LOF and ATPGobble constitute the first validated toolkit for direct manipulation of the energy state in eukaryotic cells, which will enable investigation of causal relationships between energy state and phenotypes.

Beyond activation of ATP-producing pathways and decrease in ATP/ADP ratio, we observed several other effects that have been observed previously in states involving energy depletion or AMPK activation (Cartee, 2015; McLeod & Proud, 2002; Wu et al., 2013). First, we observed activation of AMPK–an established regulator of cellular response to low energy states. AMPK activation by ATPGobble led to inhibition of *de novo* lipogenesis through phosphorylation of ACC (Figure 2), activation of glucose uptake through translocation of GLUT transporters to plasma membrane through phosphorylation of TXNIP and TBC1D4 (Figure 6), and inhibition of translation through phosphorylation of eEF2K and eEF2 (Figure 7). Second we observed inactivation of PDK–an established mediator of high energy state in mitochondria. Inactivation of PDK led to dephosphorylation and activation of PDH (Figure 6) that likely contributed to doubling of respiration rate (Figure 1). Together, these results further validate ATPGobble as a tool for direct manipulation of cellular energy state.

To showcase the value of ATPGobble as a tool, we have performed systematic proteomics and phosphoproteomics analysis. We observed large scale perturbations of protein levels and phosphorylation states with hundreds of significant changes (Figures 4 and 5). Some of the changes can be attributed to known effects of AMPK and PDK activity, which are directly regulated by ATP/ADP ratio, while many other effects do not appear to have known connections to cellular energy state and may represent either indirect effects (e.g., results of cell growth slowdown) or previously unappreciated energy sensing pathways. Some of the notable changes that we observed included concerted downregulation of proteasome core subunits (Figure 4), which may be related to the slowdown of translation and protein turnover that we observed (Figure 7), upregulation of NBPF proteins (Figure 3A) that contain DUF1220/Olduvai domain with a poorly understood function that have been implicated in human brain development and experience a rapid human lineage-specific increase in copy number, and numerous changes in cytoskeleton proteins (Figures 4, 5, 8). Deep mechanistic follow up on each of those changes is beyond the scope of this tool development study, but we hope that these results will enable domain experts to use ATPGobble to study the response of relevant biological processes to low energy state conditions.

We anticipate that ATPGobble will be useful in a variety of applications in the future. First, ATPGobble can be used to study the cellular response to low energy state in tissue culture. We have shown that ATPGobble leads to activation of energy-producing and inhibition of energy-consuming pathways through a combination of mass action, allosteric regulation, posttranslational modification and gene expression but many of the mechanistic details of how it happens remain to be elucidated. Second, we expect that ATPGobble may be useful in deciphering which phenotypes of exercise and calorie restriction are caused by perturbation of cellular energy state vs other effects. For example, ATPGobble could be expressed in muscle and the effects of this perturbation could be compared to effects of exercise. In the past, other GEMMs such as LbNOX have been successfully expressed in living animals (Levine et al., 2021; Yadav et al., 2025), suggesting that this is a plausible application.

In summary, we have developed and validated a genetically encoded tool for direct manipulation of cellular energy state. A large body of evidence connects cellular energy state with changes in physiology and disease and we hope that this reagent will be useful in future studies establishing a causal connection between cellular energy state and phenotypes.

## Supporting information

Attachment 1

Attachment 2

Attachment 3

Attachment 4

## RESOURCE AVAILABILITY

### Lead contact

Requests for further information and resources should be directed to Professor Denis V. Titov.

### Materials availability

Vectors used in this study will be deposited on Addgene by the date of final paper publication.

### Data and code availability

The mass spectrometry proteomics data associated with Figures 4 and 5 have been deposited to the ProteomeXchange Consortium via the PRIDE (Perez-Riverol et al., 2025) partner repository with the dataset identifier PXD066979. This paper does not report original code. Any additional information required to reanalyze the data reported in this paper is available from the lead contact upon request.

## ACKNOWLEDGEMENTS

We thank the members of the Titov Lab for their valuable comments on the manuscript. Research reported in this publication was supported by the National Institute of General Medical Sciences of the National Institutes of Health under award numbers DP2 GM132933 and R35 GM152114 to D.V.T. and Thanks to Scandinavia and American Scandinavian Foundation scholarships 2021–2022 to E.E.K.

We thank Dr. Gabriela Grigorean for performing the proteomics sample injection, the LC-MS/MS method writing and running in the Proteomics Core Facility of the Genome Center, University of California, Davis. The Bruker timsTOF HT LC/MS system was supported by the Howard Hughes Medical Institute, Investigator Award for Dr. Neal Hunter, UC Davis.

## AUTHOR CONTRIBUTIONS

Conceptualization, A.E.E. and D.V.T.; methodology, A.E.E, M.M.K., and D.V.T.; investigation, A.E.E. and D.V.T.; data curation, A.E.E., and D.V.T.; visualization, A.E.E. and D.V.T.; funding acquisition, D.V.T.; writing – original draft, A.E.E. and D.V.T.; writing – review & editing, A.E.E., and D.V.T.

## METHODS

### Plasmid generation

Sequences coding for atpA, atpG, and atpD subunits of *E. coli* F1 ATPase were codon-optimized using ThermoFisher codon optimizer tool and cloned into the pLVX-TetOne-Puro backbone (Takara Bio #631849). For atpA and atpD subunits, the puromycin resistance element was replaced with a blasticidin (pENTR-eGFP-attP(bxb)-*BsdR, a gift from Richard Davis – Addgene plasmid # 183751 ; http://n2t.net/addgene:183751 ; RRID:Addgene_183751) and zeocin (pUC57-GSX2-tTA-BleoR, a gift from Elena Cattaneo – Addgene plasmid #161748; http://n2t.net/addgene:161748; RRID:Addgene_161748) element respectively. As a negative catalytically inactive control, the plasmid carrying atpD subunit carrying a K155Q substitution was generated using a NEB site-directed mutagenesis kit (NEB #E0554S). Sequences coding for PercevalHR and cpmVenus were amplified from PercevalHR-FUGW (Addgene #49083) (Tantama et al., 2013) and cloned into pLVX-EF1A-Tet3G (Takara Bio #631359) instead of Tet3G. The substitutions were made using NEB HiFi DNA assembly kit and manual (NEB #E2621), with the primer design for the insert amplification done in the Benchling assembly wizard, except that for the insertions targeting the multicloning site, the backbone was cut using EcoRI and BamHI (NEB) instead of amplified. This resulted in the generation of pLVX-TetOne-bsd-atpA, pLVX-TetOne-bleo-atpD, pLVX-TetOne-bleo-atpD-K155Q and pLVX-TetOne-puro-atpG (Attachments 6–9 respectively).

### Lentivirus preparation and concentration

To generate lentiviral particles, 293T cells were seeded at 50,000 cells per cm2 and 200 uL medium (DMEM, 10% FBS), and transfected 24 hours with fresh medium, 1 ug pPAX, 1 ug plasmid of interest and 100 ng pMD per million cells, using X-tremeGENE™ 9 DNA Transfection Reagent (Roche #XTG9-RO) according to the manufacturer’s instructions, with the subsequent 2-day incubation and medium collection. Lentivirus was concentrated using the protocol provided by the University of Texas MD Anderson Cancer Center Functional Genomics Core (URL: https://www.mdanderson.org/documents/core-facilities/Functional Genomics Core/Homemade 4fold lentivirus concentrator.pdf, accessed on 11/13/2024): Briefly, the medium was centrifuged at 500g for 5 minutes, one fourth volume of the concentrator (40% PEG-8000, 1.2M NaCl PBS, pH 7.0—7.2) was added, mixed carefully, and left with agitation overnight at +4 C. The precipitate was centrifuged at 1,600 g for 1 hour at +4 C, supernatant removed, the pellet resuspended in the medium relevant to the target cell line supplemented with 8 ug/mL polybrene and the mixture was used immediately.

### Cell generation and culture

hTERT-RPE1 cells (ATCC #CRL-4000) were obtained from UC Berkeley Cell Culture Facility and maintained using DMEM/F12 (Gibco #12500062) with 10% FBS (Seradigm #89510-186) and 0.01 mg/mL hygromycin (Gibco #10687010). For short-term experiments requiring specialized media, DMEM was used instead of DMEM/F12. The target cells (hTERT-RPE1) were seeded at density of 5,000/cm2; infected 24 hours later with a medium change, containing the final 8 ug/mL of polybrene and titered concentrations of freshly concentrated lentivirus; after 8 hours of the virus treatment, medium was changed to regular maintenance medium, and relevant antibiotics (5 ug/mL blasticidin, 100 ng/mL zeocin, 10 mg/mL puromycin, 500 ug/mL geneticin) were introduced 40 hours later. Selection was assessed using parental cells as control and drugs used in all subsequent cultures.

### Seahorse assay

30,000 RPE1 cells per well were seeded 16 hours before the induction in 200 uL, then 800 uL medium added 4 hours later. The cells were induced with 300 ng/mL doxycycline for 24 hours. A Cell Mito Stress Test was run on Seahorse XFe24 using the flux assay kit (Agilent #102340-100) according to manufacturer’s instructions as follows: the cells were rinsed with 1 mL 5 mM HEPES, 10 % FBS DMEM, and left to incubate with 1 mL 5 mM HEPES, 10 % FBS DMEM at +37 C (no CO2) for an hour; 1uM oligomycin, 2.5 uM FCCP, and 1 uM antimycin was injected in 20X stocks into ports A, B, and C, respectively with the omission of rotenone to minimize side effects The measurement protocol on Wave was used with the default settings. Cells were trypsinized in 1 mL 0.25% trypsin with no rinse, pooled for each treatment group and counted using Beckman Coulter Z1 counter for normalization.

### Metabolite extraction and mass spectrometry

380,000 hTERT-RPE1 cells were seeded per dish on 6-cm dishes, 6 dishes per treatment group, next day treated with doxycycline or vehicle, and harvested 24 hours after the induction. Metabolites were extracted using 40% methanol 40% acetonitrile 20% water and 0.4% v/v formic acid. Control dishes designated for cell counting were quantified, and the rest other dishes harvested. The extraction buffer was added promptly and immediately after the removal of medium, on dry ice at an estimated ratio of 1 or 2 mL cells/mL. Each dish was harvested on its own to ensure adherence to the protocol, incubated for 5 minutes on dry ice, centrifuged at 17,000 rpm at +4C, supernatant preserved and stored at -80C.

A triple quadrupole mass spectrometer (Agilent 6430 QQQ) equipped with electrospray ionization was coupled to the HPLC system to perform targeted metabolomics via dynamic multiple reaction monitoring (dMRM). The dMRM method utilized both positive and negative ionization mode, where metabolite m/z and retention times optimized and validated using an in-house metabolite library. QQQ source parameters were held constant at: gas, 350C at 11 L/minute; nebulizer, 25 psi; capillary, 3000V (both negative and positive). Agilent MassHunter (version 10.1) was used for data acquisition, and Skyline (MacCoss Lab version 24.1.0.214) was used for analysis.

### Western blot

390,000 hTERT-RPE1 cells were seeded on 6-cm dishes, treated with doxycycline or vehicle, rinsed with PBS and lysed in 190 uL 250 uL 1% SDS Lysis buffer (50 mM Tris + 1% SDS, pH 8) with Complete Mini protease inhibitor following the manufacturer’s protocol (ThermoFisher C10102). Protein concentrations were determined using BCA and 35–40 ug protein was loaded for SDS-PAGE. Antibodies were as follows, at 1:10,000 dilution unless mentioned otherwise: calnexin (Abcam ab22595, 1:10,000), beta-actin (CST 4970L), phospho-ACC S79 (CST 11818S), phospho-AMPK T172 (CST 2535S), ACC (CST 3676S), AMPK (CST 5831S), EEF2 (CST 2332S), phospho-EEF2 T56 (2331S), phospho-PDHA S232 (CST 15289S), phospho-PDHA S293 (CST 31866S), TXNIP (CST 14715S), pan-NBPF (Thermo Fisher PA5-101712, 1:500), PDHA (CST 3205S). For phosphorylated targets, the membranes were blocked in 5% BSA-TBST, for total protein targets in 5% milk-TBST, for 1 h at room temperature, after which the primary incubation was down with 1% BSA-TBST or 1% milk-TBST respectively. The secondary antibody was anti-rabbit (CST 7074S, 1:20,000) in 1% BSA-TBST or 1% milk-TBST respectively, for 45 min at room temperature, with three 10-minute TBST washes before and after. Imaging was performed using Super Signal West femto sensitivity substrate (Thermo Scientific 34096) and images analysed using ImageJ. Phospho-targets were normalized to actin, then to their respective total-protein targets normalized to actin, after which each value was normalized to the experiment average.

### Translation assay

For translation assay, hTERT-RPE1 cells were seeded on 10 cm dishes as follows: LOF cells, three dishes 125K cells each. ATPGobble cells, three dishes 125K cells each and 1e6 cells on one dish for a 4-day induction. Dishes were induced with 300 ng/mL doxycycline for 1 or 4 days or treated with vehicle (1:166 DMSO in MilliQ water). Medium was changed every day to minimize the acidification caused by the ATPGobble activity. On the day of the sample collection, the dishes were washed with the cysteine and methionine-free DMEM containing 10 % FBS at -1 hour, one vehicle-treated ATPGobble dish was treated with 25 ug/mL cycloheximide at -30 minutes, and 50 uM AHA was added at 0 minutes. The samples were collected at 4 hours in 250 uL 1% SDS Lysis buffer (50 mM Tris + 1% SDS, pH 8) with Complete Mini protease inhibitor following the manufacturer’s protocol (ThermoFisher C10102). The click reaction and clean-up was performed following the manufacturer’s protocol (ThermoFisher C10276). 10 ug protein was loaded for analysis.

### PercevalHR

hTERT-RPE1 cells infected with the doxycycline-inducible ATPGobble or LOF constructs, as well as cpmVenus or PercevalHR, were seeded at 150K per well on a 12-well plate (Cellvis #P12-1.5H-N), induced, and the emission was imaged at 520 nm on Zeiss LSM88 using 405 nm and 488 nm excitation wavelengths, which correlate with the ADP and ATP-bound form of PercevalHR respectively. 4×4 averaging was applied to both channels to improve the signal to noise ratio of the third ratio channel. The third ratio channel was created using the calculator function in ZEN, with the formula Channel 1 / Channel 2 * 200; the multiplier was used to improve visibility without adjusting screen settings. As a positive control for a drop in ATP/ADP ratio, the vehicle wells were supplemented with 50 mM 2-deoxyglucose and 1 uM antimycin, treated for 10–15 minutes, and imaged again. One data point represents one experiment where the mean intensities from channel 3 are averaged from 5 original images. The illustration was created by saving the representative original channel 3 images in a 16-color lookup table in ImageJ. The imaging experiments were done at, and under the guidance of Biological Imaging Facility at Berkeley.

### Cell viability assay

400 RPE1 cells/well were seeded on clear-bottom, black-wall 96-well plates (Corning #3904), and induced with 300 ng/mL doxycycline (or water as vehicle control) 24 hours later, using phenol-free DMEM/F12 with glutamine, sodium pyruvate and 10% FBS. One plate was taken for imaging immediately after the induction, and then every 24 hours. For imaging, Hoechst and SYTOX Green were added to the final concentration of 2 ug/mL and 1 uM respectively, and imaging of each well in its entirety was done on Cytation 1, with 5% CO2 and at 37 C, 1 hour after the stain addition. The objects in channels acquired using DAPI and GFP cubes were counted using Gen5 3.05. As a positive control for SytoxGreen, hTERT-RPE1 cells were treated with 50 mM 2-deoxyglucose and 1 uM antimycin.

### Cell line growth comparison

The infection protocol was repeated as described above to generate a duplicate of the hTERT-RPE1, while infecting the additional cell lines outlined in Table S1. Since hTERT-RPE1 reaches confluence at a relatively small cell number, for the other cell lines, 50,000 cells per well were seeded. After a two-week drug selection, cell growth was quantified using the above cell viability assay, with cell seeding numbers modified as shown in Table S1.

### Cell growth assay for mitomycin validation

hTERT-RPE1-LOF and ATPGobble cells were grown and treated as described in the cell viability assay with the addition of 700 ng/mL mitomycin. Instead of live imaging, the plates were fixed with 4% PFA, after which they were stained with 1 ug/mL Hoechst for 30 minutes room temperature, stored at +4C and imaged the same day, with the nuclei counted using Gen5 3.05 software.

### Proteomics cell culture

The experiment was prepared in three batches, with two replicates in each batch, resulting in a dataset of 16 total protein samples and 16 enriched phosphopeptide samples. For each replicate, RPE1 cell lines expressing ATPGobble constructs as described were seeded at 2e6 per dish on six 15 cm dishes for the ATPGobble doxycycline group, and three 15 cm dishes for each other group. In addition, one cell-counting dish was seeded for each condition. The expression was induced 24 hours after the seeding and samples collected 24 hours after the induction.

### Proteomics sample preparation

A protocol provided by Cell Signaling Technology was adapted (CST #14989) with several modifications. Briefly, the cell dishes were rinsed with cold PBS and lysed in 1 mL lysis buffer (9 M urea, 20 mM HEPES pH 8.0, 2X Phosphatase Inhibitor Cocktail) per 5e6 cells, passed 10 times through a 25G needle, then cleared at 17,000 x g for 15 min at room temperature. Concentrations were measured using BCA (ThermoFisher #23225), amounts and volumes equalized, reduced in 4.5 mM DTT for 60 min, alkylated in 10 mM iodoacetamide for 15 minutes in the dark, diluted 4.5-fold with 20 mM HEPES pH 8.0, then digested in 37.5 ug trypsin per mg protein at RT on a shaker overnight. The peptides were desalted using C18 reversed-phase Sep-Pak^™^ columns from Waters (#WAT054955), eluted in 1.5 mL 0.1% TFA, 50% acetonitrile and dried in a SpeedVac in 100 uL and 1.4 mL, after which the 100 uL pellet was used for the total proteomics analysis and the 1.4 mL pellet was used for the phosphopeptide enrichment and analysis.

### Phosphoproteomics sample preparation

Phosphopeptides were enriched using TiO2 Phosphopeptide Enrichment Kit (ThermoFisher A32993) following the kit instructions.

### Mass spectrometry run

Five hundred ng total peptide was loaded onto a disposable Evotip C18 trap column (Evosep Biosytems, Denmark) and subjected to nanoLC on a Evosep One instrument (Evosep Biosystems). Tips were eluted directly onto a PepSep analytical column, dimensions: 150umx25cm C18 column (PepSep, Denmark) with 1.5 μm particle size (100 Å pores) (Bruker Daltronics). Mobile phases A and B were water with 0.1% formic acid (v/v) and 80/20/0.1% ACN/water/formic acid (v/v/vol), respectively. The standard pre-set method of 30samples-per-day (44minute run) was used. The mass spectrometry was done on a hybrid trapped ion mobility spectrometry-quadrupole time of flight mass spectrometer (timsTOF HT, (Bruker Daltonics, Bremen, Germany), operated in PASEF mode. The acquisition scheme used for DIA consisted of 36 precursor windows at width of 25m/z, over the mass range 300-1200 Dalton. The TIMS scans layer the doubly and triply charged peptides.

### Proteomics data analysis

The raw data was analyzed using DIA-NN 2.1.0 with a human fasta library from Uniprot version UP000005640_9606 that featured additional sequences for the ATPGobble constructs, with parameters set as follows: --qvalue 0.01 --matrices --min-corr 2.0 --corr-diff 1.0 time-corr-only --extracted-ms1 --met-excision --cut K*,R* --missed-cleavages 1 -- unimod4 --var-mods 1 --var-mod UniMod:21,79.966331,STY --mass-acc 15 --mass-acc-ms1 15 --individual-mass-acc --individual-windows --peptidoforms --reanalyse --rt-profiling. cRAP sequences were removed, and the values from pg_matrix for total proteomics and report.phosphosites_99.tsv for phosphoproteomics were normalized by column intensity. The data was imputed using a tail-end imputation method based on a set of conditions shown in Table S2. After the tail-end imputation, the rows with more than 7 missing values in either cell line were deleted, and the rest of the missing values were imputed using a pseudorandom number generator within one row standard deviation of the row median. The essential code used to carry out imputation and filtering is shown in Attachments 9 and 10.

### Scratch assay

150,000 hTERT-RPE1 cells were seeded on a 12-well plate and treated with doxycycline or vehicle, with the addition of 700 ng/mL mitomycin at the same time. 24 hours after the induction, the cells were scratched and medium changed with the experiment conditions maintained. The scratch crosses were imaged using a BioTek Cytation 3 plate reader at 0, 12 and 24 hours after the scratch, and the bare area was quantified using ImageJ.

---

**Figure S1.**
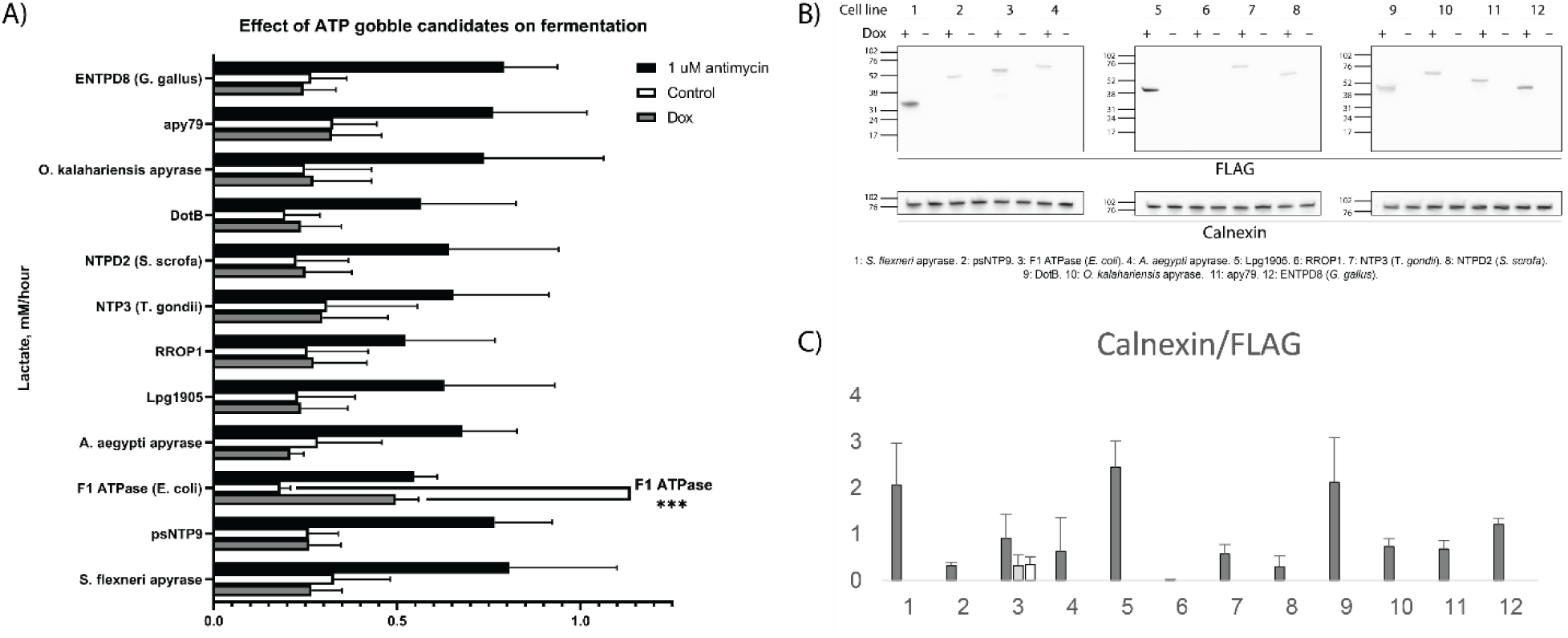
Screening for ATPGobble candidates. 1: RSFA (*S. flexneri* apyrase). 2: psNTP9. 3: F1-ATPase. 4: *A. aegypti* apyrase. 5: Lpg1905. 6: RROP1. 7: NTP3. 8: NTPD2. 9: DotB. 10: *O. kalahariensis* apyrase. 11: apy79. 12: NTPD8. A) Lactate assay showing no ATPGobble activity for most candidates except F1 ATPase from E. coli. B) Western blot showing the expression levels of the FLAG-tagged candidates used for the lactate assay and C) Western blot quantification (n = 3). Experiment means + standard deviation of experiment means.

**Figure S2.**
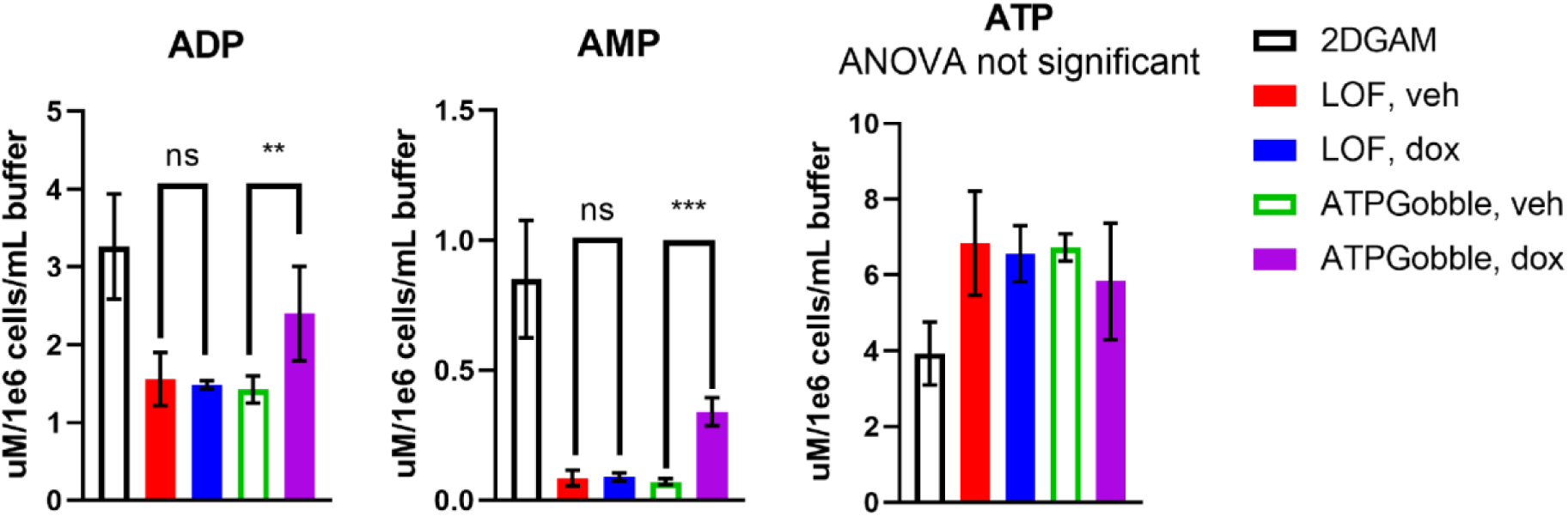
ATPGobble expression in hTERT-RPE1 increases the concentrations of ADP and AMP, while ATP is maintained at a stable level. 3 LC/MS experiments 5 biological replicates each. Repeated measurements ANOVA followed by the main row effect test with Tukey’s adjustment for multiple comparisons. ns > 0.05, * < 0.05, ** < 0.01 *** < 0.001. dox: doxycycline, veh: vehicle. Experiment means ± standard deviation of experiment means.

**Figure S3.**
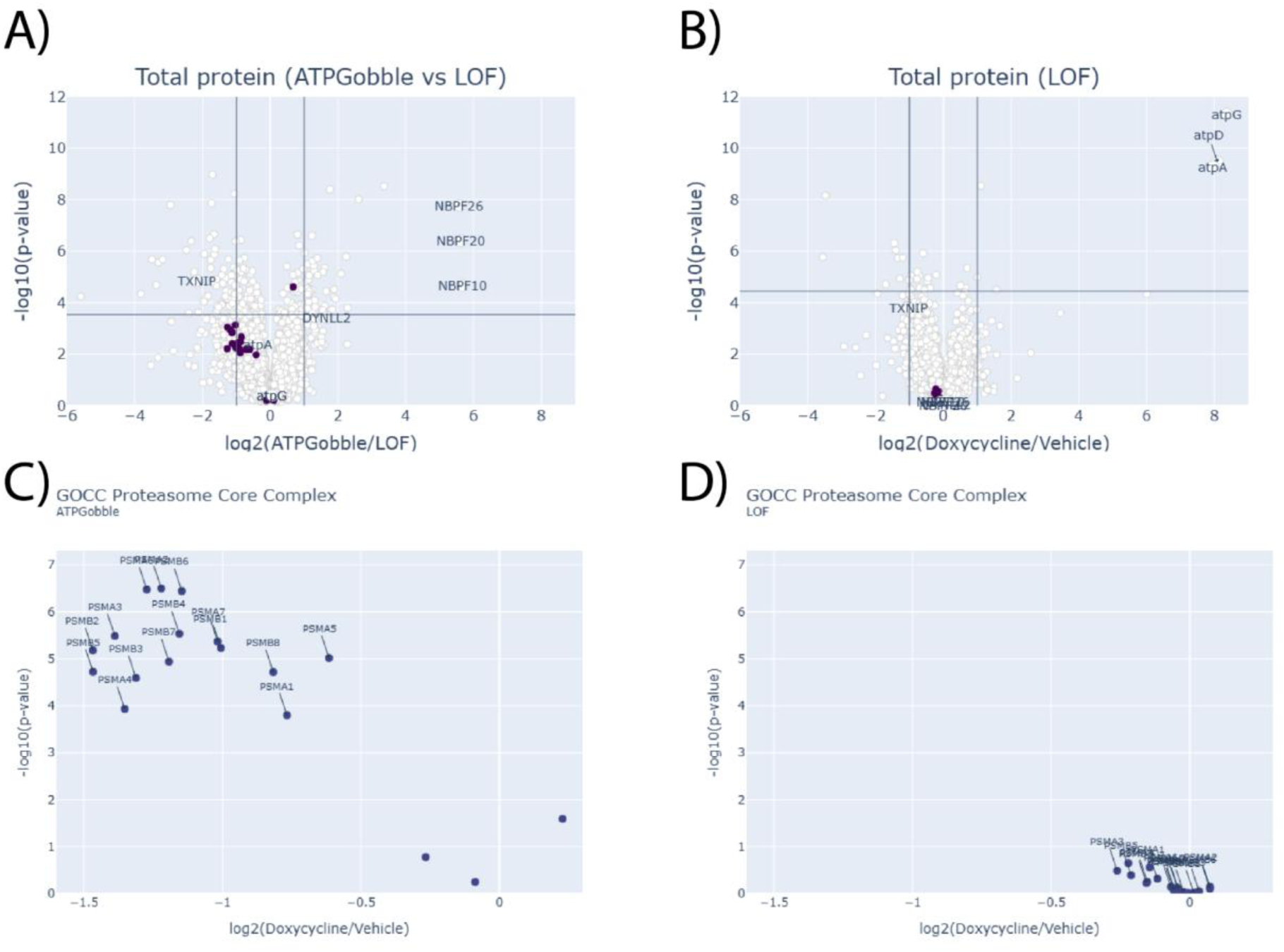
Alternative comparisons of total proteomics data and breakdown of proteasome core data set. A) Volcano plot comparing doxycycline treated hTERT-RPE1-LOF and ATPGobble cells. B) Volcano plot comparing vehicle vs doxycycline treated hTERT-RPE1-LOF cells. Volcano plots showing the changes in the expression of proteasome core complex proteins after vehicle vs doxycycline treatment in hTERT-RPE1-ATPGobble (C) vs LOF (D) cells.

**Figure S4.**
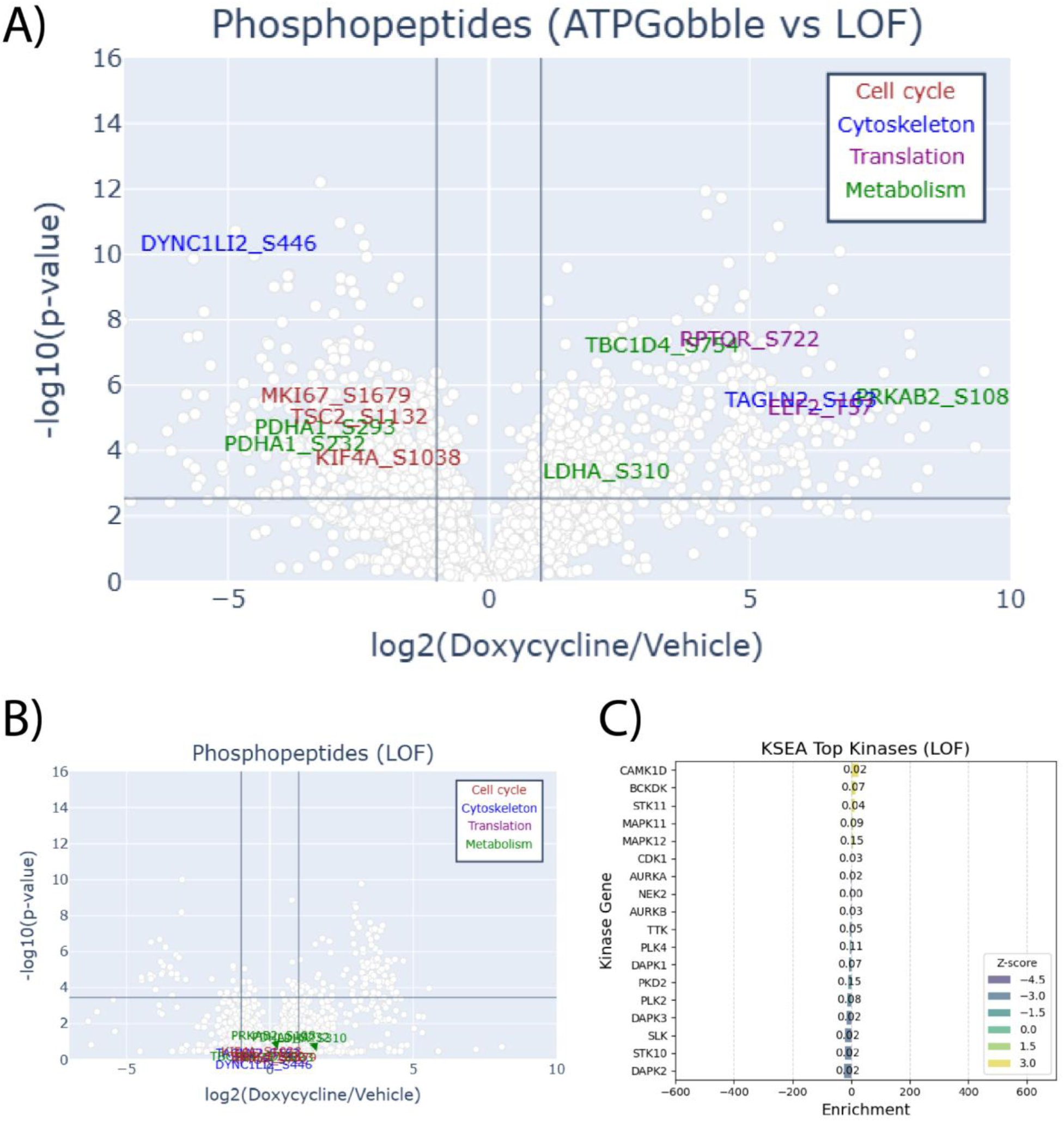
ATPGobble activity results in phosphoproteomic changes – supplementary analyses. A) A volcano plot comparing phosphopeptides from hTERT-RPE1-LOF and hTERT-RPE-ATPGobble cells after doxycycline treatment. B) A volcano plot comparing hTERT-RPE1-LOF cells treated with vehicle vs doxycycline. The targets and the axis ranges shown in A and B are the same as in Figure 5A. C) KSEA of hTERT-RPE1-LOF cells after vehicle vs doxycycline treatment. The X axis range is the same as in Figure 5C.

**Figure S5.**
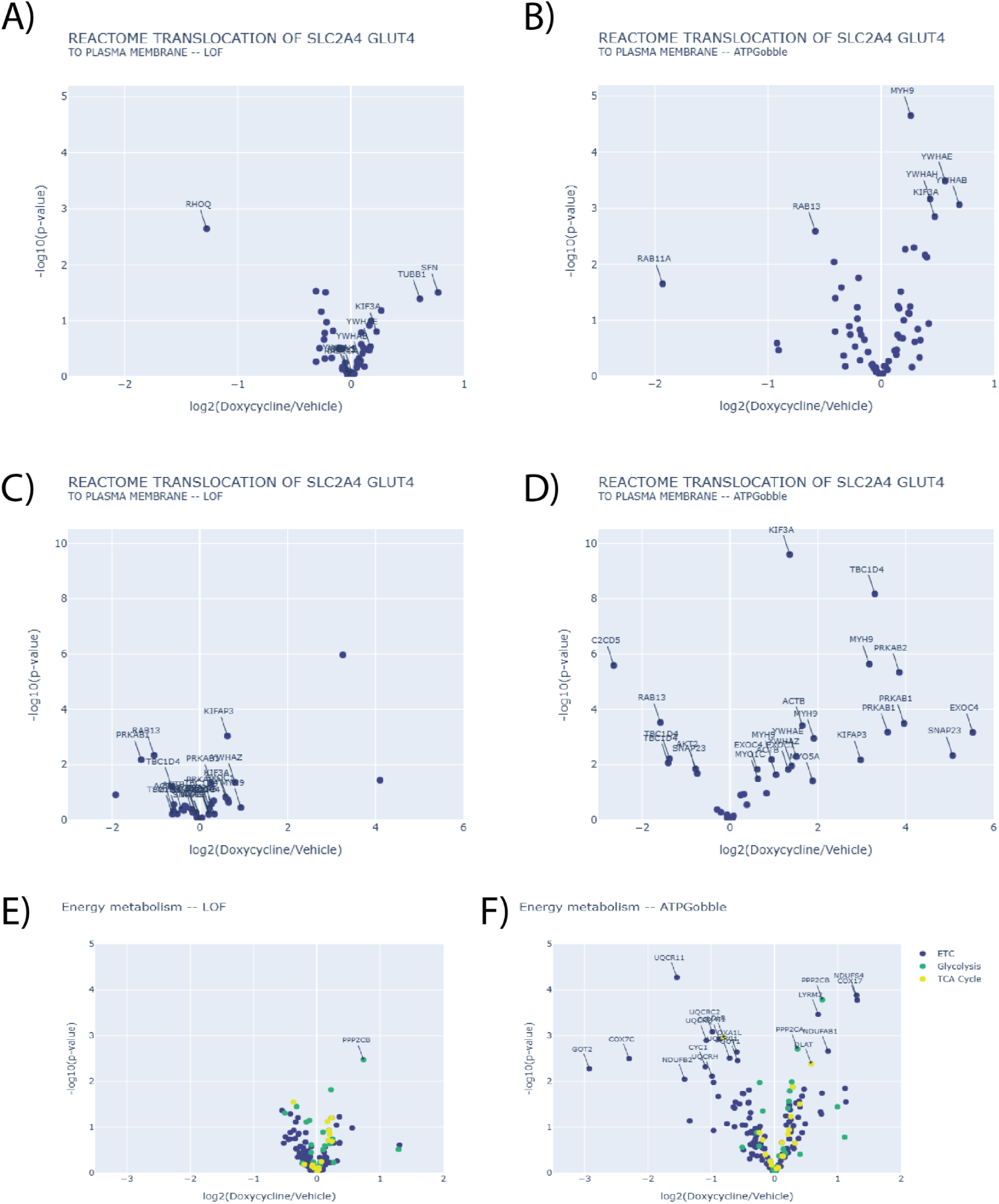
Changes in energy-producing activity in ATPGobble-expressing cells. Total-protein changes in proteins regulating the translocation of GLUT4 (coded by *SLC2A4*) to the membrane in hTERT-RPE1-LOF (A) and hTERT-RPE1-ATPGobble (B) cells after vehicle vs doxycycline treatment. Phosphopeptide changes of the same gene set in hTERT-RPE1-LOF (C) and hTERT-RPE1-ATPGobble (D) cells after vehicle vs doxycycline treatment. Changes in genes coding for ETC, glycolysis and TCA cycle machinery, LOF (E) and ATPGobble (F).

**Figure S6.**
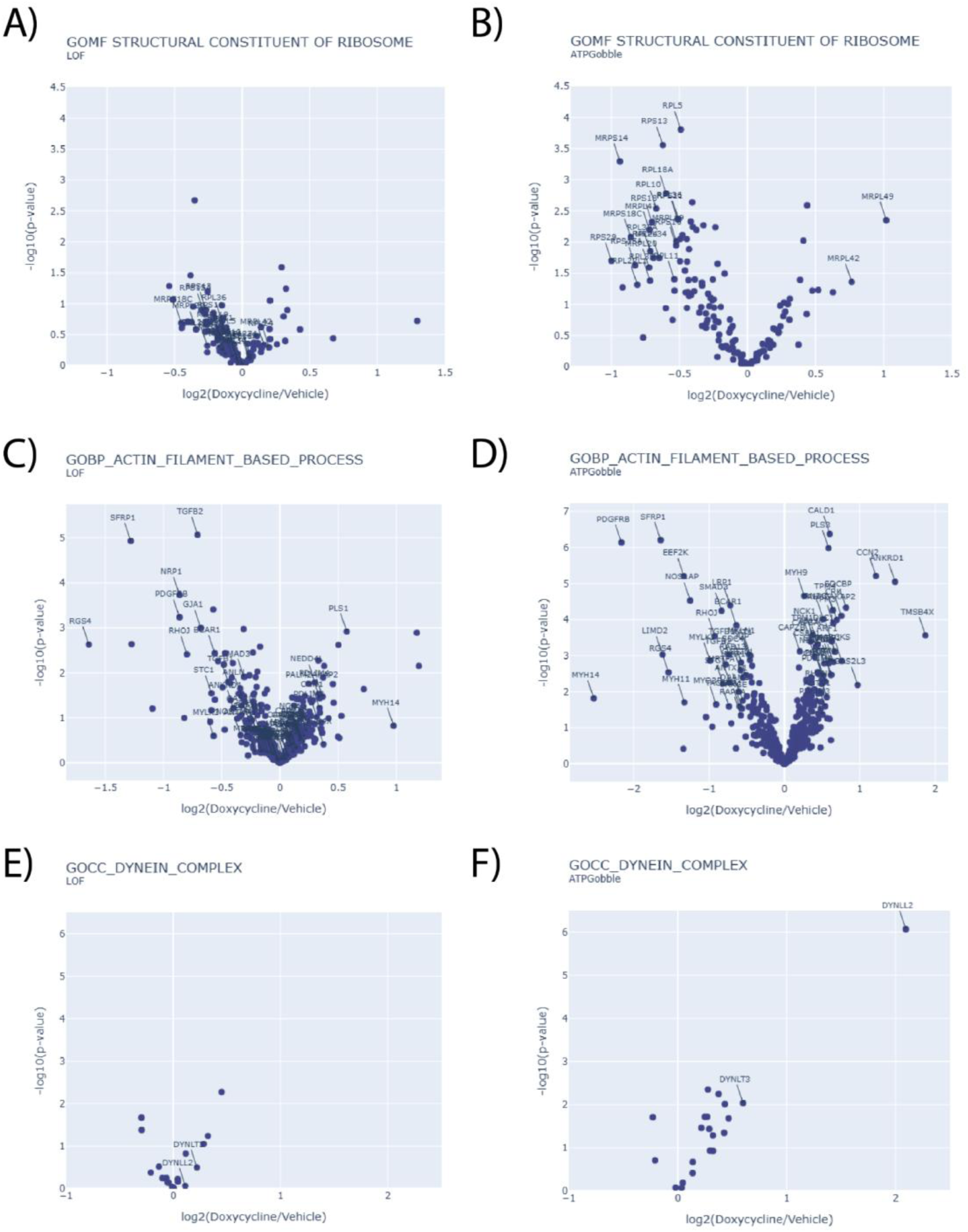
Changes in energy-consuming activity in cells expressing ATPGobble. Changes in GOMF STRUCTURAL CONSTITUENT OF RIBOSOME proteins in hTERT-RPE1-LOF (A) and hTERT-RPE1-ATPGobble (B) cells after vehicle vs doxycycline treatment. Changes in GOBP ACTIN FILAMENT BASED PROCESS proteins in hTERT-RPE1-LOF (C) and hTERT-RPE1-ATPGobble (D) cells after vehicle vs doxycycline treatment. Changes in GOCC DYNEIN COMPLEX proteins in hTERT-RPE1-LOF (E) and hTERT-RPE1-ATPGobble (F) cells after vehicle vs doxycycline treatment.

**Table S1.**
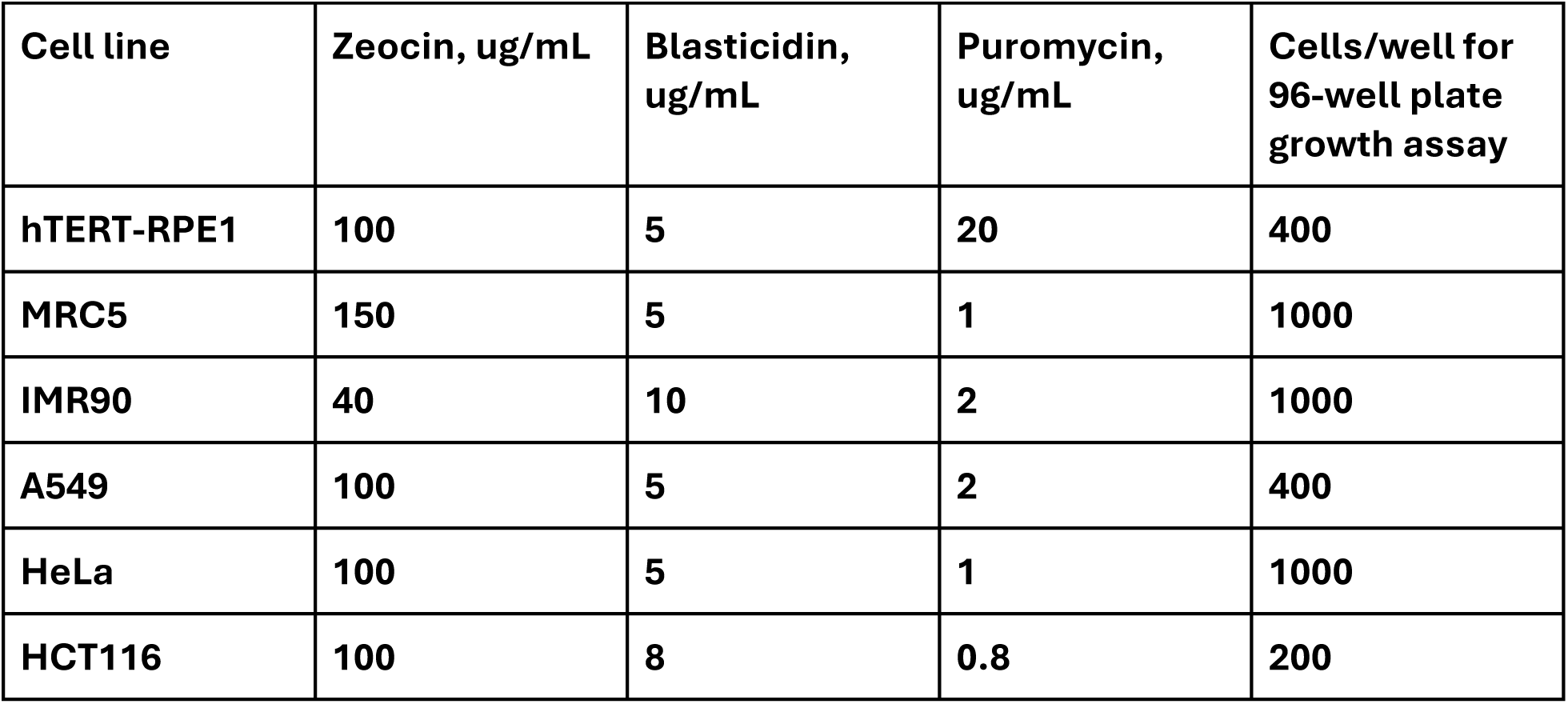
Cell line generation and seeding density for cell growth comparison.

**Table S2.** Summary of steps for the tail-end imputation.

| **Table S2. Summary of steps for the tail-end imputation.** |  |  |
| --- | --- | --- |
| **Data set** | **Condition** | **Impute with** |
| Total proteomics<br>(report.pg_matrix.tsv) | All values in dox/veh group and no values in the opposite group. | minimum*0.2*(1-abs(random.gauss(0, 0.5)))<br>(try again if the generated number below 0) |
| Total proteomics<br>(report.pg_matrix.tsv) | 4 or more missing values in dox/veh group and at most 1 missing value in the opposing group. | minimum*(1-abs(random.gauss(0, 0.25))) (try again if the generated number below 0) |
| Phosphopeptides<br>(report.phosphosites_99.tsv) | At least 3 non-NaN values in dox/veh group and all NaN values in the opposing group | minimum*0.2*(1-abs(random.gauss(0, 0.5)))<br>(try again if the generated number below 0) |
| Phosphopeptides<br>(report.phosphosites_99.tsv) | At least 4 non-NaN values in dox/veh group and less than 3 non-NaN values in the opposing group. | minimum*(1-abs(random.gauss(0, 0.25))) (try again if the generated number below 0) |

## References

Ashcroft, F. M., & Rorsman, P. (1989). Electrophysiology of the pancreatic beta-cell. Progress in Biophysics and Molecular Biology, 54(2), 87–143. 10.1016/0079-6107(89)90013-8

Atkinson, D. E. (1968). The energy charge of the adenylate pool as a regulatory parameter. Interaction with feedback modifiers. Biochemistry, 7(11), 4030–4034. 10.1021/bi00851a033

Biondi, R. M., Komander, D., Thomas, C. C., Lizcano, J. M., Deak, M., Alessi, D. R., & van Aalten, D. M. F. (2002). High resolution crystal structure of the human PDK1 catalytic domain defines the regulatory phosphopeptide docking site. The EMBO Journal, 21(16), 4219–4228. 10.1093/emboj/cdf437

Boecker, S., Slaviero, G., Schramm, T., Szymanski, W., Steuer, R., Link, H., & Klamt, S. (2021). Deciphering the physiological response of Escherichia coli under high ATP demand. Molecular Systems Biology, 17(12), e10504. 10.15252/msb.202110504

Browne, G. J., Finn, S. G., & Proud, C. G. (2004). Stimulation of the AMP-activated protein kinase leads to activation of eukaryotic elongation factor 2 kinase and to its phosphorylation at a novel site, serine 398. The Journal of Biological Chemistry, 279(13), 12220–12231. 10.1074/jbc.M309773200

Carling, D., Zammit, V. A., & Hardie, D. G. (1987). A common bicyclic protein kinase cascade inactivates the regulatory enzymes of fatty acid and cholesterol biosynthesis. FEBS Letters, 223(2), 217–222. 10.1016/0014-5793(87)80292-2

Cartee, G. D. (2015). AMPK-TBC1D4–Dependent Mechanism for Increasing Insulin Sensitivity of Skeletal Muscle. Diabetes, 64(6), 1901–1903. 10.2337/db15-0010

Caskey, C. T., Ashton, D. M., & Wyngaarden, J. B. (1964). THE ENZYMOLOGY OF FEEDBACK INHIBITION OF GLUTAMINE PHOSPHORIBOSYLPYROPHOSPHATE AMIDOTRANSFERASE BY PURINE RIBONUCLEOTIDES. The Journal of Biological Chemistry, 239, 2570–2579.

Choe, M., & Titov, D. V. (2022). Genetically encoded tools for measuring and manipulating metabolism. Nature Chemical Biology, 18(5), 451–460. 10.1038/s41589-022-01012-8

Demichev, V., Messner, C. B., Vernardis, S. I., Lilley, K. S., & Ralser, M. (2020). DIA-NN: Neural networks and interference correction enable deep proteome coverage in high throughput. Nature Methods, 17(1), 41–44. 10.1038/s41592-019-0638-x

Denton, R. M., Randle, P. J., Bridges, B. J., Cooper, R. H., Kerbey, A. L., Pask, H. T., Severson, D. L., Stansbie, D., & Whitehouse, S. (1975). Regulation of mammalian pyruvate dehydrogenase. Molecular and Cellular Biochemistry, 9(1), 27–53. 10.1007/BF01731731

Eickelschulte, S., Hartwig, S., Leiser, B., Lehr, S., Joschko, V., Chokkalingam, M., Chadt, A., & Al-Hasani, H. (2021). AKT/AMPK-mediated phosphorylation of TBC1D4 disrupts the interaction with insulin-regulated aminopeptidase. The Journal of Biological Chemistry, 296, 100637. 10.1016/j.jbc.2021.100637

Gabriel, J. L., Milner, R., & Plaut, G. W. (1985). Inhibition and activation of bovine heart NAD-specific isocitrate dehydrogenase by ATP. Archives of Biochemistry and Biophysics, 240(1), 128–134. 10.1016/0003-9861(85)90015-3

Glunčić, M., Vlahović, I., Rosandić, M., & Paar, V. (2024). Neuroblastoma Breakpoint Family 3mer Higher Order Repeats/Olduvai Triplet Pattern in the Complete Genome of Human and Nonhuman Primates and Relation to Cognitive Capacity. Genes, 15(12), 1598. 10.3390/genes15121598

Hucho, F., Randall, D. D., Roche, T. E., Burgett, M. W., Pelley, J. W., & Reed, L. J. (1972). α-Keto acid dehydrogenase complexes: XVII. Kinetic and regulatory properties of pyruvate dehydrogenase kinase and pyruvate dehydrogenase phosphatase from bovine kidney and heart. Archives of Biochemistry and Biophysics, 151(1), 328–340. 10.1016/0003-9861(72)90504-8

Ibrahim, A., Yucel, N., Kim, B., & Arany, Z. (2020). Local Mitochondrial ATP Production Regulates Endothelial Fatty Acid Uptake and Transport. Cell Metabolism, 32(2), 309. 10.1016/j.cmet.2020.05.018

Koebmann, B. J., Westerhoff, H. V., Snoep, J. L., Nilsson, D., & Jensen, P. R. (2002). The glycolytic flux in Escherichia coli is controlled by the demand for ATP. Journal of Bacteriology, 184(14), 3909–3916. 10.1128/JB.184.14.3909-3916.2002

Lawlis, V. B., & Roche, T. E. (1981). Regulation of bovine kidney alpha-ketoglutarate dehydrogenase complex by calcium ion and adenine nucleotides. Effects on S0.5 for alpha-ketoglutarate. *Biochemistry*, *20*(9), 2512–2518. 10.1021/bi00512a023

Levine, D. C., Kuo, H.-Y., Hong, H.-K., Cedernaes, J., Hepler, C., Wright, A. G., Sommars, M. A., Kobayashi, Y., Marcheva, B., Gao, P., Ilkayeva, O. R., Omura, C., Ramsey, K. M., Newgard, C. B., Barish, G. D., Peek, C. B., Chandel, N. S., Mrksich, M., & Bass, J. (2021). NADH inhibition of SIRT1 links energy state to transcription during time-restricted feeding. Nature Metabolism, 3(12), 1621–1632. 10.1038/s42255-021-00498-1

Lowry, O. H., & Passonneau, J. V. (1966). Kinetic Evidence for Multiple Binding Sites on Phosphofructokinase. Journal of Biological Chemistry, 241(10), 2268–2279. 10.1016/S0021-9258(18)96616-0

McLeod, L. E., & Proud, C. G. (2002). ATP depletion increases phosphorylation of elongation factor eEF2 in adult cardiomyocytes independently of inhibition of mTOR signalling. FEBS Letters, 531(3), 448–452. 10.1016/S0014-5793(02)03582-2

Ohta, Y., Taguchi, A., Matsumura, T., Nakabayashi, H., Akiyama, M., Yamamoto, K., Fujimoto, R., Suetomi, R., Yanai, A., Shinoda, K., & Tanizawa, Y. (2017). Clock Gene Dysregulation Induced by Chronic ER Stress Disrupts β-cell Function. EBioMedicine, 18, 146. 10.1016/j.ebiom.2017.03.040

Perez-Riverol, Y., Bandla, C., Kundu, D. J., Kamatchinathan, S., Bai, J., Hewapathirana, S., John, N. S., Prakash, A., Walzer, M., Wang, S., & Vizcaíno, J. A. (2025). The PRIDE database at 20 years: 2025 update. Nucleic Acids Research, 53(D1), D543–D553. 10.1093/nar/gkae1011

Pratt, M. L., & Roche, T. E. (1979). Mechanism of pyruvate inhibition of kidney pyruvate dehydrogenasea kinase and synergistic inhibition by pyruvate and ADP. Journal of Biological Chemistry, 254(15), 7191–7196. 10.1016/S0021-9258(18)50303-3

Rubink, D. S., & Winder, W. W. (2005). Effect of phosphorylation by AMP-activated protein kinase on palmitoyl-CoA inhibition of skeletal muscle acetyl-CoA carboxylase. Journal of Applied Physiology (Bethesda, Md.: 1985), 98(4), 1221–1227. 10.1152/japplphysiol.00621.2004

Sakai, K., Kawashima, S., Suzuki, K., & Imahori, K. (1987). Affinity labeling of the allosteric site of fructose 1,6-bisphosphatase with an AMP analog. Journal of Biochemistry, 102(2), 377–384. 10.1093/oxfordjournals.jbchem.a122064

Schulze, J. O., Saladino, G., Busschots, K., Neimanis, S., Süß, E., Odadzic, D., Zeuzem, S., Hindie, V., Herbrand, A. K., Lisa, M.-N., Alzari, P. M., Gervasio, F. L., & Biondi, R. M. (2016a). Bidirectional Allosteric Communication between the ATP-Binding Site and the Regulatory PIF Pocket in PDK1 Protein Kinase. Cell Chemical Biology, 23(10), 1193–1205. 10.1016/j.chembiol.2016.06.017

Schulze, J. O., Saladino, G., Busschots, K., Neimanis, S., Süß, E., Odadzic, D., Zeuzem, S., Hindie, V., Herbrand, A. K., Lisa, M.-N., Alzari, P. M., Gervasio, F. L., & Biondi, R. M. (2016b). Bidirectional Allosteric Communication between the ATP-Binding Site and the Regulatory PIF Pocket in PDK1 Protein Kinase. Cell Chemical Biology, 23(10), 1193–1205. 10.1016/j.chembiol.2016.06.017

Senior, A. E., & al-Shawi, M. K. (1992). Further examination of seventeen mutations in Escherichia coli F1-ATPase beta-subunit. Journal of Biological Chemistry, 267(30), 21471–21478. 10.1016/S0021-9258(19)36633-5

Treebak, J. T., Taylor, E. B., Witczak, C. A., An, D., Toyoda, T., Koh, H.-J., Xie, J., Feener, E. P., Wojtaszewski, J. F. P., Hirshman, M. F., & Goodyear, L. J. (2010). Identification of a novel phosphorylation site on TBC1D4 regulated by AMP-activated protein kinase in skeletal muscle. American Journal of Physiology - Cell Physiology, 298(2), C377–C385. 10.1152/ajpcell.00297.2009

Ureta, T. (1976). The allosteric regulation of hexokinase C from amphibian liver. The Journal of Biological Chemistry, 251(16), 5035–5042.

Vishnu, N., Hamilton, A., Bagge, A., Wernersson, A., Cowan, E., Barnard, H., Sancak, Y., Kamer, K. J., Spégel, P., Fex, M., Tengholm, A., Mootha, V. K., Nicholls, D. G., & Mulder, H. (2021). Mitochondrial clearance of calcium facilitated by MICU2 controls insulin secretion. Molecular Metabolism, 51, 101239. 10.1016/j.molmet.2021.101239

Waite, K. A., De-La Mota-Peynado, A., Vontz, G., & Roelofs, J. (2016). Starvation Induces Proteasome Autophagy with Different Pathways for Core and Regulatory Particles. The Journal of Biological Chemistry, 291(7), 3239–3253. 10.1074/jbc.M115.699124

Weber, G. (1969). Regulation of pyruvate kinase. Advances in Enzyme Regulation, 7, 15–40. 10.1016/0065-2571(69)90007-7

Winder, W. W., & Hardie, D. G. (1996). Inactivation of acetyl-CoA carboxylase and activation of AMP-activated protein kinase in muscle during exercise. American Journal of Physiology-Endocrinology and Metabolism, 270(2), E299–E304. 10.1152/ajpendo.1996.270.2.E299

Woods, A., Vertommen, D., Neumann, D., Türk, R., Bayliss, J., Schlattner, U., Wallimann, T., Carling, D., & Rider, M. H. (2003). Identification of Phosphorylation Sites in AMP-activated Protein Kinase (AMPK) for Upstream AMPK Kinases and Study of Their Roles by Site-directed Mutagenesis*. Journal of Biological Chemistry, 278(31), 28434–28442. 10.1074/jbc.M303946200

Wu, N., Zheng, B., Shaywitz, A., Dagon, Y., Tower, C., Bellinger, G., Shen, C.-H., Wen, J., Asara, J., McGraw, T. E., Kahn, B. B., & Cantley, L. C. (2013). AMPK-dependent degradation of TXNIP upon energy stress leads to enhanced glucose uptake via GLUT1. Molecular Cell, 49(6), 1167–1175. 10.1016/j.molcel.2013.01.035

Yadav, S., Pan, X., Li, S., Martin, P. L., Hoang, N., Chen, K., Karhadkar, A., Malhotra, J., Zuckerman, A. L., Munan, S., Klose, M. K., Wang, L., Cracan, V., & Parkhitko, A. A. (2025). Tissue-specific modulation of NADH consumption as an anti-aging intervention in Drosophila. bioRxiv: The Preprint Server for Biology, 2025.01.06.631511. 10.1101/2025.01.06.631511

